# Optogenetic generation of leader cells reveals a force-velocity relation for collective cell migration

**DOI:** 10.1101/2024.01.23.576733

**Authors:** Leone Rossetti, Steffen Grosser, Juan Francisco Abenza, Léo Valon, Pere Roca-Cusachs, Ricard Alert, Xavier Trepat

## Abstract

The front of migratory cellular clusters during development, wound healing and cancer invasion is typically populated with highly protrusive cells that are called leader cells. Leader cells are thought to physically pull and direct their cohort of followers, but how leaders and followers are mechanically organized to migrate collectively remains controversial. One possibility is that the autonomous local action of a leader cell is sufficient to drive migration of the group. Yet another possibility is that a global mechanical organization is required for the group to move cohesively. Here we show that the effectiveness of leader-follower organization is proportional to the asymmetry of traction and tension within the cellular cluster. By combining hydrogel micropatterning and optogenetic activation of Rac1, we locally generate highly protrusive leaders at the edge of minimal cell groups. We find that the induced leader can robustly drag one follower but is generally unable to direct larger groups. By measuring traction forces and tension propagation in groups of increasing size, we establish a quantitative relationship between group velocity and the asymmetry of the traction and tension profiles. We propose a model of the motile cluster as an active polar fluid that explains this force-velocity relationship in terms of asymmetries in the distribution of active tractions. Our results challenge the notion of autonomous leader cells by showing that collective cell migration requires a global mechanical organization within the cluster.

## Introduction

Collective motion is a recurrent property of groups of self-propelled agents, including macromolecular assemblies, human crowds or robots swarms^1–3^. The coordination and function of these groups can emerge from interactions between identical constituents^4,5^. However, group coordination is often mediated by specialized agents^6^. Small numbers of such agents can either disrupt or enhance group dynamics^7–9^, like in the case of animal groups guided by specialized individuals that act as leaders^10,11^. Similarly, collectively migrating cells during development, wound healing or collective cancer invasion are thought to be guided by a subgroup of leader cells^12–20^.

Leader cells are found at the front edge of migrating cell groups, have a protrusive and polarized phenotype and are identified by the activity of specific signalling pathways^14,17,21–26^. Conversely, the remaining cells are termed follower cells and are thought to be mechanically pulled and guided by their leaders through a direct physical connection^15,22,27–29^. This description appears to capture many *in vitro* and *in vivo* phenomena. Epithelial cell sheets migrate by projecting multi-cellular outgrowths, with protrusive cells at their front^13,23,25^. Similarly, during branching morphogenesis, long cell strands are tipped by lamellipodium-generating cells^30,31^. Border cells are small clusters of ∼8 cells that migrate during Drosophila oogenesis and have a polarized and protrusive cell at their front^27,32^. In some invasive tumours, thin strands of cells project out from a cluster following a single cancer cell or a cancer-associated fibroblast^26,33–35^.

Despite this extensive phenomenology, the fundamental mechanical organization that enables leader-follower coordination is still unclear. Two main scenarios are possible. The first one is that the local mechanical action of an autonomous leader is sufficient to drive the migration of a group, regardless of the behaviour of its followers. A second scenario is that a global organization of forces within the group is needed for collective migration. Addressing this long-standing problem requires a direct measurement of the relationship between collective cell velocity and the underlying spatial distribution of forces, but such a force-velocity relationship has not been reported. To fill this gap, here we used optogenetics to induce leader cells in minimal groups of controlled size. We show that generation of a leader is insufficient to drive the migration of groups larger than two cells. To understand this behaviour, we performed a systematic study of the mechanical conditions that enable collective cell migration in clusters of increasing cell number. This analysis revealed that, for every cluster size, collective cell velocity requires an asymmetric distribution of traction forces on a multicellular scale. A model of the cell cluster as an active polar fluid establishes the relationship between the velocity of a cell group and the asymmetries of the underlying traction force field.

## Results

### Generating minimal leader-follower systems using optogenetic control of Rac1 activity

A hallmark of leader cells is lamellipodium formation by the activation of the small Rho GTPase Rac1 at their leading edge^12,15,36^. We reasoned that we could use optogenetics to locally activate Rac1 in cell groups, thus creating leader cells on demand^32,37,38^. To achieve this, we generated stable lines of MDCK cells (optoMDCK-Rac1) expressing two constructs: CIBN-GFP-CAAX and TIAM-CRY2-mCherry; the first is targeted to the plasma membrane, while the second one is cytosolic and carries the catalytic domain of Tiam1, an activator of Rac1. Upon illumination with blue light, the two constructs bind with high affinity, localizing Tiam1 at the membrane^32,37,39–41^ (Fig. 1a). Previous work has shown that Rac1 is activated within the illuminated region^37^. As expected, the photoactivation of optoMDCK-Rac1 cells results in the formation of a lamellipodium (Fig. 1b) and of focal adhesions (Extended Data Fig. 1a).

**Figure 1.**
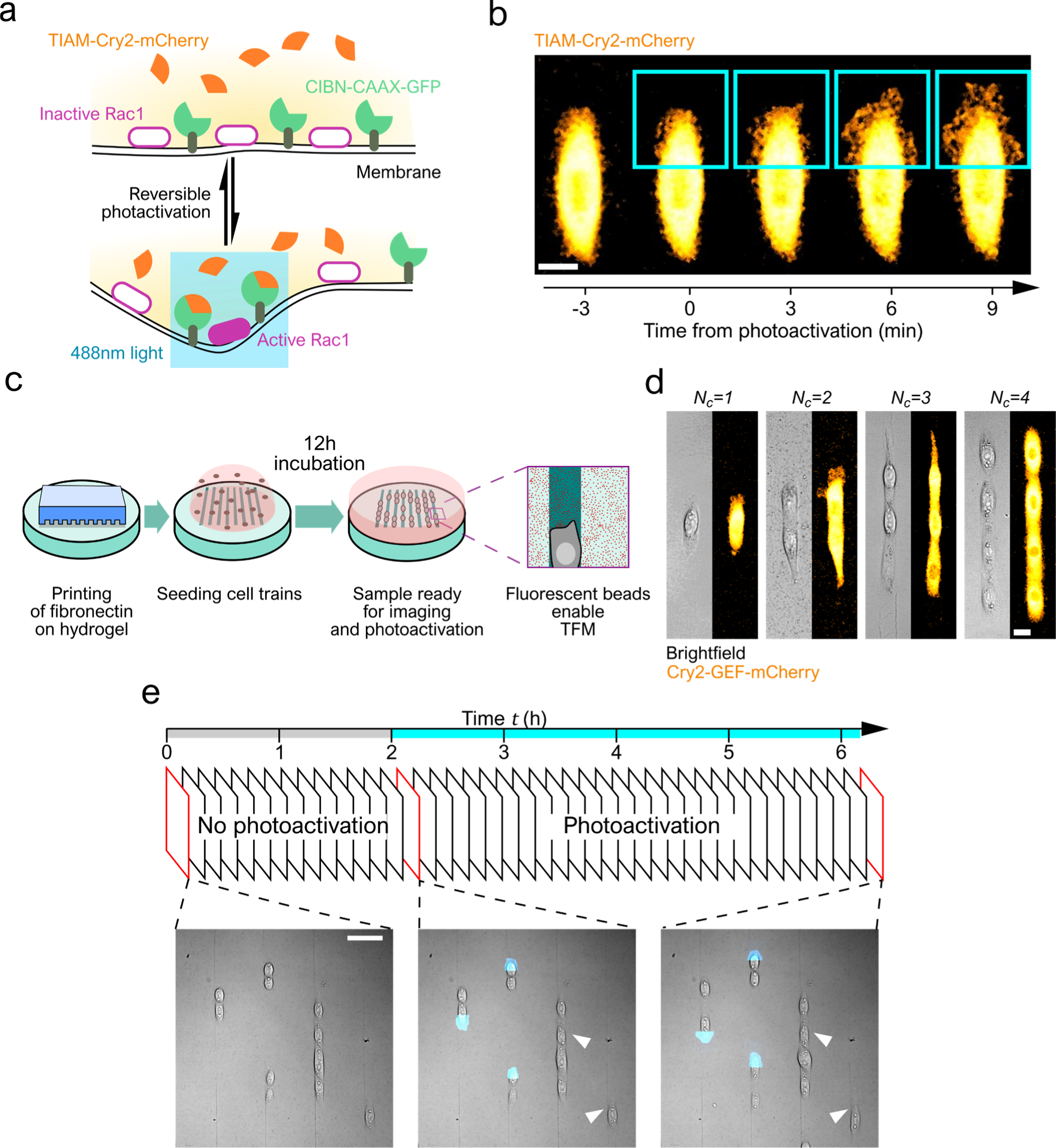
Optogenetic control of lamellipodium formation in cell trains. **(a)** Scheme of the optogenetic system to control lamellipodium formation. OptoMDCK-Rac1 express two constructs: an activator of Rac1 fused to Cry2 and membrane-bound CIBN. Upon blue-light illumination, Cry2 binds to CIBN locally activating Rac1 and causing lamellipodium growth. **(b)** Effect of photoactivation on a cell. 488 nm light is applied to the blue region every 3 minutes, inducing lamellipodium formation (scale bar 20 µm). **(c)** Scheme of sample preparation: microcontact printing of polyacrylamide hydrogels (E=18 kPa) with fibronectin lines (width 20 µm) and incubation with optoMDCK-Rac1 yields samples containing hundreds of cell trains of different lengths. Substrates are prepared with fluorescent beads, making them apt for traction force microscopy. **(d)** Representative microscopy images of cell trains (scale bar 20 µm). **(e)** Scheme of the experimental protocol. Fields of view containing cell trains (*N*_*c*_ = [1,4]) are imaged for 2 hours every 3 minutes without photoactivation, then a subset of cell trains is photoactivated (blue regions) at every imaging interval while other cell trains (white arrowheads) are left non-activated.

To study the mechanical coupling of leaders and followers we engineered highly reproducible minimalistic systems that captured the fundamental elements of collective motion. We used microcontact printing on polyacrylamide gels of uniform stiffness (18 kPa) to create fibronectin-coated lines 20 µm wide and several millimetres long. We seeded optoMDCK-Rac1 cells at a low concentration so that they attached to the patterns individually and in small linear groups, which we refer to as “cell trains”, with a sparse spatial distribution (Fig. 1c). The cells were allowed to adhere on the lines for 12 hours and were maintained in media containing thymidine to halt cell division. The polyacrylamide gels were prepared containing fluorescent microspheres so that traction force microscopy (TFM) could be performed^42^.

We imaged regions of the sample containing several isolated cell trains, ranging in length from one to four cells (*N*_*c*_ = 1,2,3,4; Fig. 1d). The imaging was initially performed for two hours without photoactivation, acquiring mCherry and microsphere fluorescence. Following this baseline measurement, some cell trains were photoactivated while the others were not. In the photoactivated trains, a rectangle of blue light illumination was located at one of their free edges, inducing lamellipodia formation. The photoactivation and the imaging continued for 4-5 hours (Fig. 1e), and the position of the photoactivated regions was periodically updated to follow the movement of the cells (Fig. 2a, Supplementary Movies 1-4). At the end of each experiment, cells were detached from the gel and an image of the relaxed microspheres was acquired for TFM calculations.

**Figure 2.**
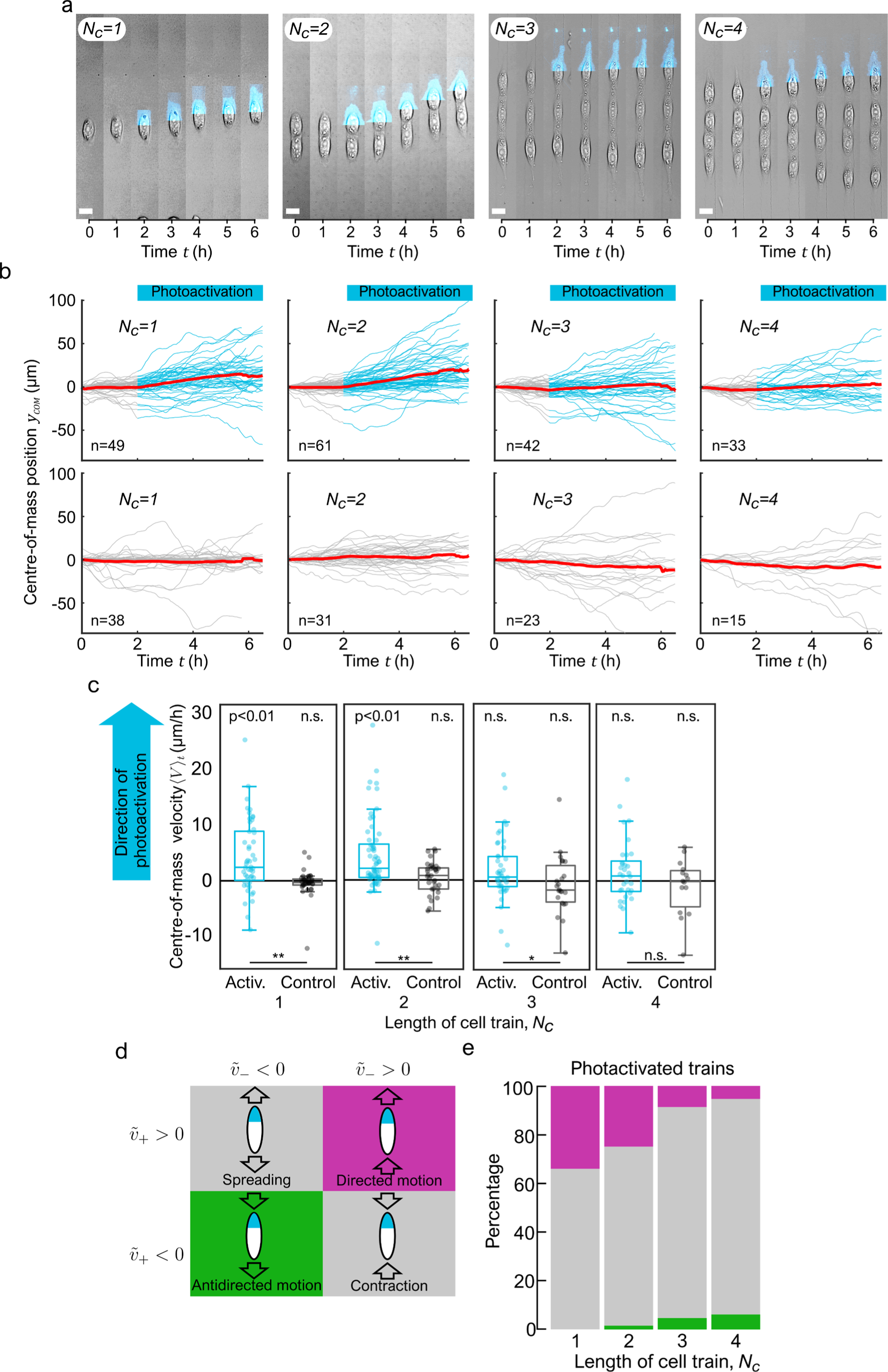
Leader cell migratory efficiency decreases with number of followers. **(a)** Representative cell trains during photoactivation of a cell edge (scale bars 20 µm). **(b)** Centre of mass trajectories for all cell trains. Top row: photoactivated trains, blue curves represent duration of photoactivation. Bottom row: control trains. Red lines are average trajectories. **(c)** Average migration velocities of the centre of mass of all photoactivated and control cell trains. Statistical significance quantified by a two-sided Wilcoxon rank sum test. For the two-sample tests ** indicates p<0.01 and * indicates p<0.05. **(d)** Types of cell train migration based on the motion of the top and bottom edges. **(e)** Motion of photoactivated cell trains according to the definitions in (d). Magenta represents the percentage of trains that undergo collective migration in the direction of the photoactivation (i.e., directed migration), green is in the opposite direction (antidirected migration). Percentages of directed migration are: 32.6% (*N*_*c*_ =, 27.9% (*N*_*c*_ = 2), 9.5% (*N*_*c*_ = 3), 6.0% (*N*_*c*_ = 4), total sample sizes are n=49, n=61, n=42, n=33, respectively.

### A photoactivated cell can only lead the migration of one follower

Prior to any analysis, we reoriented images so that the photoactivated edge of the cell trains was always towards *y* > 0. We used a custom algorithm to segment the cell trains and measure their position, for both the photoactivated and the non-photoactivated ones. With these data we calculated *V*(*t*), the centre-of-mass velocity of the cell trains. In the non-photoactivated cases, we calculated each cell train’s average centre-of-mass velocity ⟨*V*⟩_*t*_ over the duration of the full experiment, while in the photoactivated cases we computed ⟨*V*⟩_*t*_ from one hour after photoactivation until the end of the experiment, to exclude any transient behaviour. As expected, regardless of the cell number *N*_*c*_, we observed that non-photoactivated cell trains have average velocities symmetrically distributed around 0, showing no preferred direction (Fig. 2b bottom row, c). On the other hand, the velocities of photoactivated cell trains with *N*_*c*_ = 1 are biased towards the direction of the photoactivated edge and generally display larger magnitude than the corresponding non-photoactivated cases, confirming our ability to generate moving cells on demand. Cell trains with *N*_*c*_ = 2 also show motion significantly biased towards the photoactivated edge. However, this directional bias is lost for *N*_*c*_ ≥ 3, with average velocities ⟨*V*⟩_*t*_ not significantly different between photoactivated and non-photoactivated cell trains (Fig. 2b top row, c).

To investigate if the photoactivated cell was affecting the motion of the entire train, and not only of the centre-of-mass, we analysed separately the trajectories of the top and bottom edges. For each train, we considered the sequences of instantaneous velocities of top and bottom edges *v*_+_(*t*) and *v*_−_(*t*), respectively. We defined “coherent motion” of a cell-train the case in which both top edge and bottom edge velocity medians 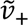 and 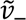 were significantly non-zero and had the same sign. We then termed “directed motion” the case with positive velocities (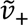 > 0, 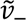 > 0, Fig. 2d), corresponding to coherent motion in the direction of photoactivation. Conversely, we categorized the opposite case as “antidirected motion” (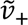 < 0, 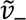 < 0). The cases that did not exhibit coherent motion were termed spreading (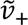 > 0, 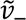 < 0) and contraction (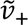 < 0, 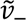 > 0).

By analysing all photoactivated cell trains with this criterion, we found that when *N*_*c*_ = 1 a third of the single-cell trains exhibit directed motion, while no single cells are moving opposite to the photoactivation (Fig. 2e). Consistent with our data on centre-of-mass velocity, this effect is rapidly lost as *N*_*c*_ grows: the percentage of cell trains with directed motion decreases, and the percentage with antidirected motion increases (Fig. 2e). For *N*_*c*_ ≥ 3 the behaviours of photoactivated cell trains are comparable to those of the non-photoactivated trains; there is no increase in directed motion due to photoactivation (Extended Data Fig. 2a).

We checked whether this behaviour was due to an inability of photoactivated cells to generate a lamellipodium when connected to more followers. To do so, we measured the extent of lamellipodium growth caused by photoactivation. We found that for all values of *N*_*c*_ the photoactivated trains display significant increases in lamellipodium size, and that the extent of this increase does not depend on the length of the train (Extended Data Fig. 1b,c). Moreover, we found that the percentage of photoactivated top edges where 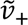 is significantly larger than 0 was >50% for all values of *N*_*c*_ (Extended Data Fig. 2b), supporting that the ability of photoactivation to induce protrusion and migration is independent of train length. We also tested whether the response of the cell trains to photoactivation was influenced by their behaviour before photoactivation. We found that nearly all trains that undergo coherent motion after photoactivation had initially been either spreading or contracting, indicating that no directedness existed prior to photoactivation (Extended Data Fig. 3).

Taken together, our experiments show that induced leader cells are not capable of driving coherent motion of a group consisting of more than one follower.

### Coherent motion requires asymmetric traction and tension fields

To identify the conditions that determine whether coherent group motion occurs or not, we turned to our measurements of the traction forces exerted by the cells on the substrate (Fig. 3a, b). Because of the linear geometry of the cell trains, we analysed *T*_*y*_, the y-component of the tractions. As expected, for *N*_*c*_ = 1 the pattern of cell tractions is that of a contractile dipole (Fig. 3b). Longer cell trains exhibit more complex patterns that are not simple superpositions of *N*_*c*_ dipoles, indicating that the cells in a train are not acting as mechanically independent units (Fig. 3b, c).

**Figure 3.**
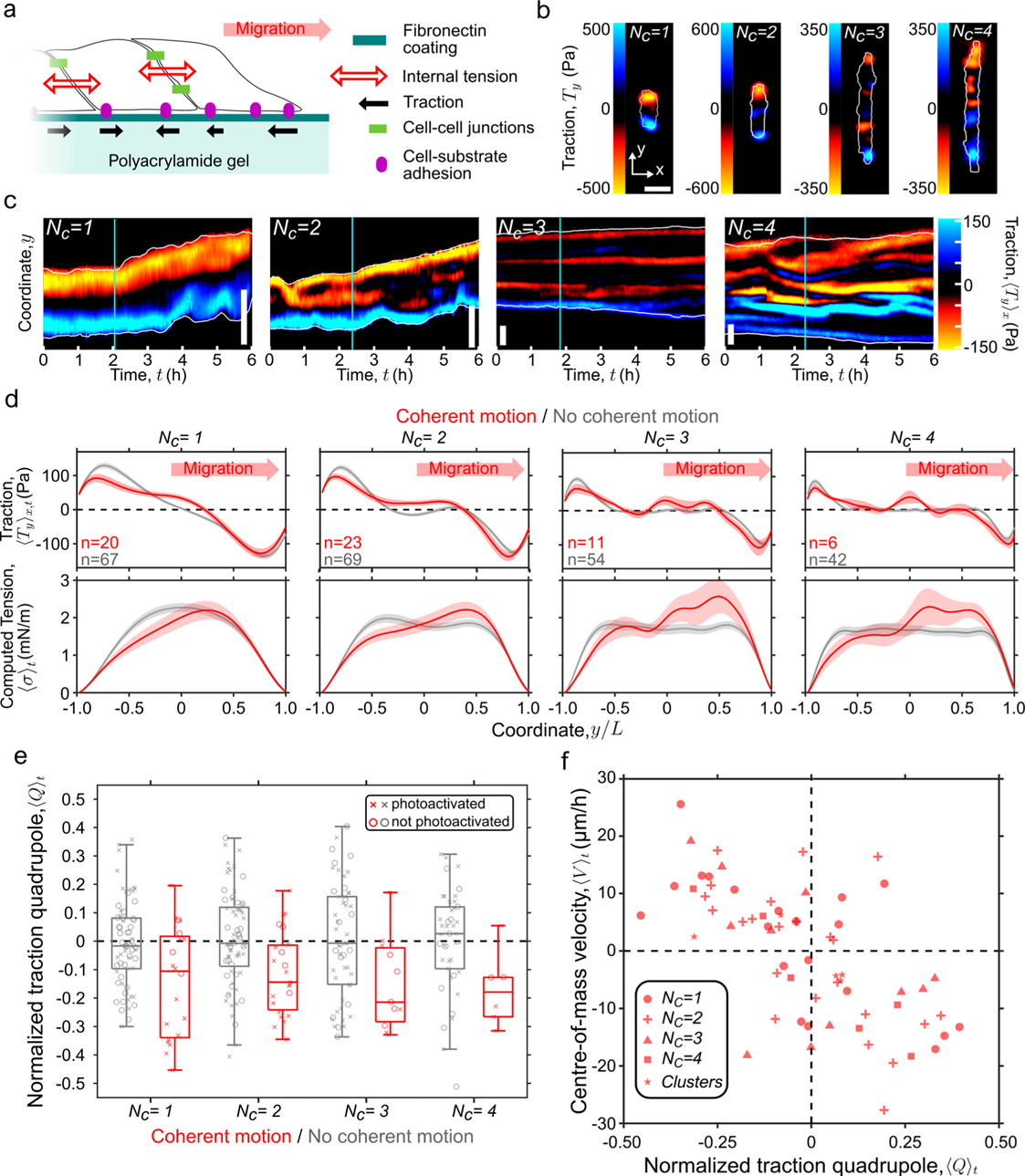
Asymmetric traction and tension profiles drive the migration of cell trains. **(a)** Scheme of cell-substrate traction and tension within a cell train. **(b)** Representative fields of the longitudinal traction forces for trains of different lengths. **(c)** Representative kymographs of the longitudinal tractions. Cyan lines mark the beginning of the photoactivation intervals, white lines are the cell edges (scale bars 50 µm). **(d)** Average profiles of longitudinal component of the traction forces (top row) and of internal tension (bottom row), for cell trains undergoing coherent motion (red) and for other trains (grey). Shaded regions along the curves show the standard error of the mean. Averages are over different cell trains; number of cell-trains *n* is indicated on the plot. **(e)** Normalized traction quadrupole of trains exhibiting coherent movement (red) and other trains (grey). **(f)** Scatter plot of average centre-of-mass velocity and normalized traction quadrupole in cell trains and clusters (see Fig. 4) exhibiting coherent motion. R=0.66, p<0.01. For clarity, in this plot the cell train velocities have not been aligned with the positive axis.

To reduce fluctuations between and within cell trains, we averaged *T*_*y*_ across the train width *x* and then computed a time-averaged traction profile 〈*T*_*y*_〉_*x,t*_ for different values of *N*_*c*_. We then compared these profiles between cell trains that did and did not move coherently. Our data shows that coherent motion occurs both in photoactivated trains (Fig. 2e) and, with a lower probability, in non-photoactivated ones (Extended Data Fig. 2a). Therefore, we binned our data in two groups, one including all coherently moving trains and another one including all non-coherently moving ones, regardless of whether the motion was directed or antidirected and whether the trains were photoactivated or non-photoactivated (see Extended Data Fig. 4 for different groupings of these data). To compare moving trains regardless of their direction of motion, we aligned the data with *y* > 0 in the direction of coherent motion. The average traction profiles of trains that do not move coherently are symmetric, with tractions of equal magnitude concentrated at the train edges (grey curves Fig. 3d, top row). With increasing train length, the tractions in the central region vanish, meaning that while edge tractions are sustained in time, central tractions are transient and tend to cancel out. By contrast, in trains undergoing coherent movement, 〈*T*_*y*_〉_*x,t*_ becomes asymmetric; the tractions at the trailing edge are lower in magnitude and extend further into the train even reaching the central region (red curves in Fig. 3d, top row).

To quantify this mechanical asymmetry, we computed the time-average of the 1D normalized traction quadrupole *Q* = (∫ *T*_*y*_ *y*^2^d*y*)/ (∫|*T*_*y*_|*y*^2^d*y*), which is the normalized second moment of the traction field along the train axis with respect to the centre of mass^43–46^. Since we are considering the coherent motion of all trains as being towards positive *y*, the quadrupole *Q* is negative for traction profiles like those of the trains undergoing coherent movement, as the integral is dominated by negative tractions concentrated at the leading edge. We found that in nearly all trains undergoing coherent movement 〈*Q*〉_*t*_ is indeed negative, while in the other trains it takes positive and negative values with equal probability (Fig. 3e).

To better understand this mechanical asymmetry, we studied the distribution of internal tension within the cell trains. To this end we calculated *σ*, the tension transmitted inside cells by the cytoskeleton and between cells by cell-cell junctions. The internal tension balances the tractions at the cell-substrate interface and in a 1D system it is given by 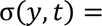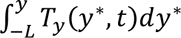, where 2*L* is the length of the cell train^29^. Analogously to our calculation of 〈*T*_*y*_〉_*x,t*_, we computed ensemble averages of *σ*. We found that trains that do not exhibit coherent movement have symmetric average tension profiles, while the trains that move coherently have an asymmetric tension profile with a broad peak located closer to the leading edge (Fig. 3d bottom row).

We then wondered if the value of 〈*Q*〉_*t*_ was related to the centre-of-mass velocity ⟨*V*⟩_*t*_. In Fig. 3a-e, we aligned all data so that the coherent motion of trains was in the direction of increasing *y*. Now, however, to better visualize the relationship between velocity and traction asymmetry and to ascertain the absence of axial bias in the experiments, we went back to considering trains moving with either positive or negative velocity, as they occurred in the experiments. Strikingly, we found that for all trains exhibiting coherent motion, ⟨*V*⟩_*t*_ and 〈*Q*〉_*t*_ are correlated. Faster trains have stronger traction asymmetries, resulting in tension that is more concentrated towards their leading edge (Fig. 3f).

Taken together, these results show that train movement is driven by the global spatial distribution of mechanical stress. Front-back asymmetries along the train are necessary for coherent motion, and stronger asymmetries drive faster motion.

### Migrating 2D clusters and monolayer fingers show mechanical asymmetries similar to 1D trains

The directed migration of 1D cell trains is common in developmental processes such as angiogenesis^47^ and branching morphogenesis^48^. This type of migration has also been observed during collective cancer invasion through small interstitial spaces^49^. In many other processes, collective cell migration involves 2D clusters or multicellular fingers protruding from a 2D cell sheet^23^. We thus asked whether the mechanical asymmetries observed in migrating 1D cell trains are also present in 2D clusters and multicellular fingers.

To study 2D cluster migration, we patterned wider fibronectin lines (50 µm across) and seeded optoMDCK-Rac1 cells so that they attached to the patterns in small clusters 2-3 cells wide (Fig. 4a-c). We selected clusters that contained 5-15 cells and photoactivated their top edge cells, applying the same experimental protocol and analysis that we used on the cell trains. Similarly to the behaviour of long trains, only a small fraction of clusters migrated coherently (∼11%). We then computed the time-averaged traction profile 〈*T*_*y*_〉_*x,t*_ for the clusters that did not move coherently and for those that did. Like in cell trains, the traction profile of the clusters that do not move coherently is symmetric (Fig. 4e, grey curves). By contrast, in clusters that move coherently, tractions at the leading edge are higher in magnitude and those at the trailing edge extend further into the cluster (Fig. 4e, red curves). Accordingly, the tension in these cell clusters has an asymmetric distribution with higher values towards the leading edge (Fig. 4g). Plotting the values of ⟨*V*⟩_*t*_ and 〈*Q*〉_*t*_ for the migrating clusters in Fig. 3f, we found that they follow a similar behavior than the cell trains.

**Figure 4.**
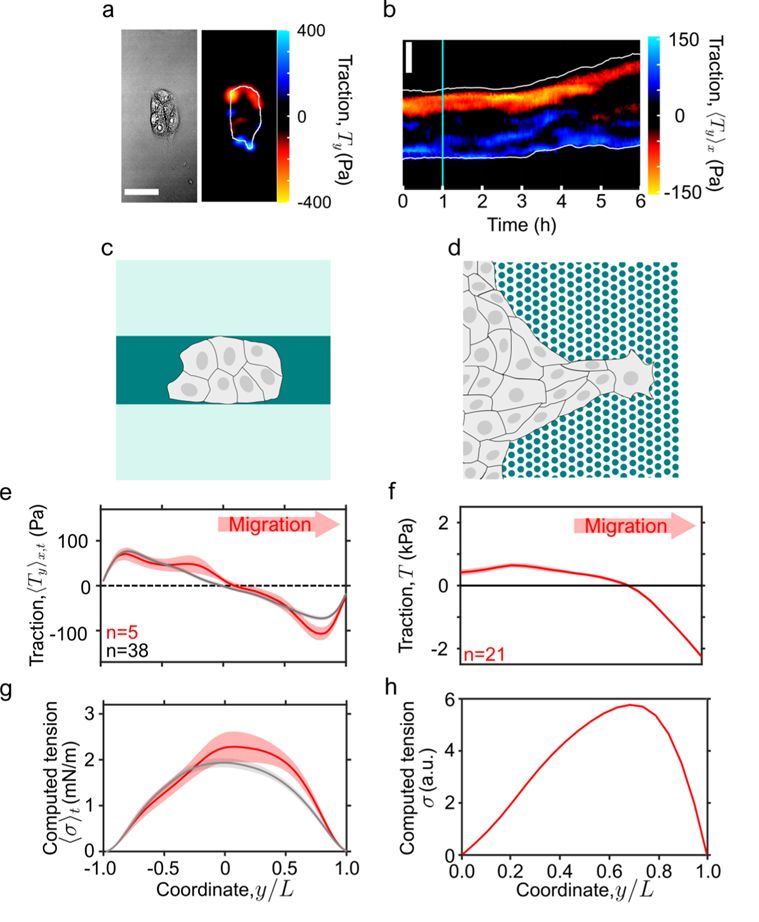
Asymmetric traction distributions appear during coherent migration of islands and invading monolayers. **(a)** Phase contrast image and longitudinal traction forces of a cell island (scale bar 50 µm). **(b)** Kymograph of the longitudinal tractions. Cyan lines mark the beginning of the photoactivation intervals (scale bar 50 µm). **(c)** Diagram of a cell island on a micropatterned fibronectin line (dark green) on a polyacrylamide gel (light green). **(d)** Diagram of a migrating finger from a monolayer edge on pillars. **(e)** Average profiles of longitudinal component of the traction forces, for cell islands undergoing coherent motion (red) and for other trains (grey). Shaded regions represent the standard error. Averages are over different cell islands; number of cell-trains *n* is indicated on the plot. **(f)** Average profile of longitudinal component of the traction forces, for migrating cell fingers. Shaded region represents the standard error of the mean. Averages are over different cell fingers; their number *n* is indicated on the plot. **(g)** Average profiles of the computed tension for cell islands undergoing coherent motion (red) and for other trains (grey). Shaded regions represent the standard error of the mean. **(h)** Average profiles of the computed tension for migrating cell fingers.

Next, we looked at the multicellular finger-like protrusions that form at the edge of an expanding epithelial monolayer. For this, we obtained data from a previous study by Reffay *et al.*, where the traction forces generated by several MDCK fingers were measured using micropillars^23^. We analysed the component of the tractions along the finger’s axis, averaged across the width of the finger analogously to how we calculated 〈*T*_*y*_〉_*x,t*_ for trains and clusters (Fig. 4d). The resulting average traction profile shows an asymmetry similar to that of coherently moving trains and clusters: tractions are higher and more localized at the leading edge than at the trailing edge (Fig. 4f). This profile results in an asymmetric tension with higher values towards the direction of migration (Fig. 4h). These findings suggest that the relationship between directed collective cell migration and tension asymmetries is a general principle that applies to both 1D and 2D systems.

### An active fluid model explains the relation between net motion and traction asymmetries

So far, our results show that an autonomous leader cell is not sufficient to drive coherent group motion. We instead find that coherent motion requires a supracellular traction and tension asymmetry, and that there is a correlation between the extent of this asymmetry and the migration velocity. To understand how supracellular tension distributions might drive collective migration, we developed an active-matter model. We modelled a cell train as a one-dimensional compressible active fluid that exerts tractions on the underlying substrate^50^. Force balance reads *δ*_*y*_*σ* = *T*, where *σ* = *σδ*_*y*_*v* is the internal tension, with *σ* being the effective viscosity and *v* the velocity field. *T* is the total traction that results from viscous drag on the substrate, *ξv*, and from myosin-generated cytoskeletal forces termed active tractions, *T*_*a*_.

Thus, the total traction is

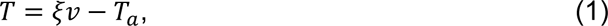

and the force balance condition is given by

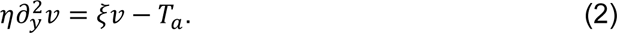

When a cell polarizes, it develops spatial asymmetries in cytoskeletal force generation. To study how these asymmetries drive motion, we took an active traction profile given by

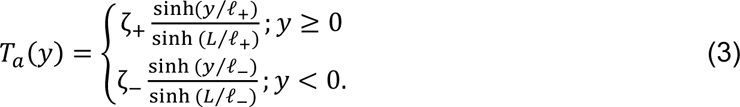

Here, ζ is the maximal active traction, which is exerted at the train edges *y* = ±*L*, and *λ* is the length determining its decay toward the train interior (Fig. 5a). Equation (3) generalizes the active traction profiles obtained in previous models of cell monolayers^50–53^ to now account for spatial asymmetries. Taking different values of and *λ* at each train edge, we captured two sources of asymmetry, namely in the magnitude and in the penetration of active tractions. Below, we analyse how these sources of asymmetry shape the total traction and tension profiles, and hence how they drive train migration.

**Figure 5.**
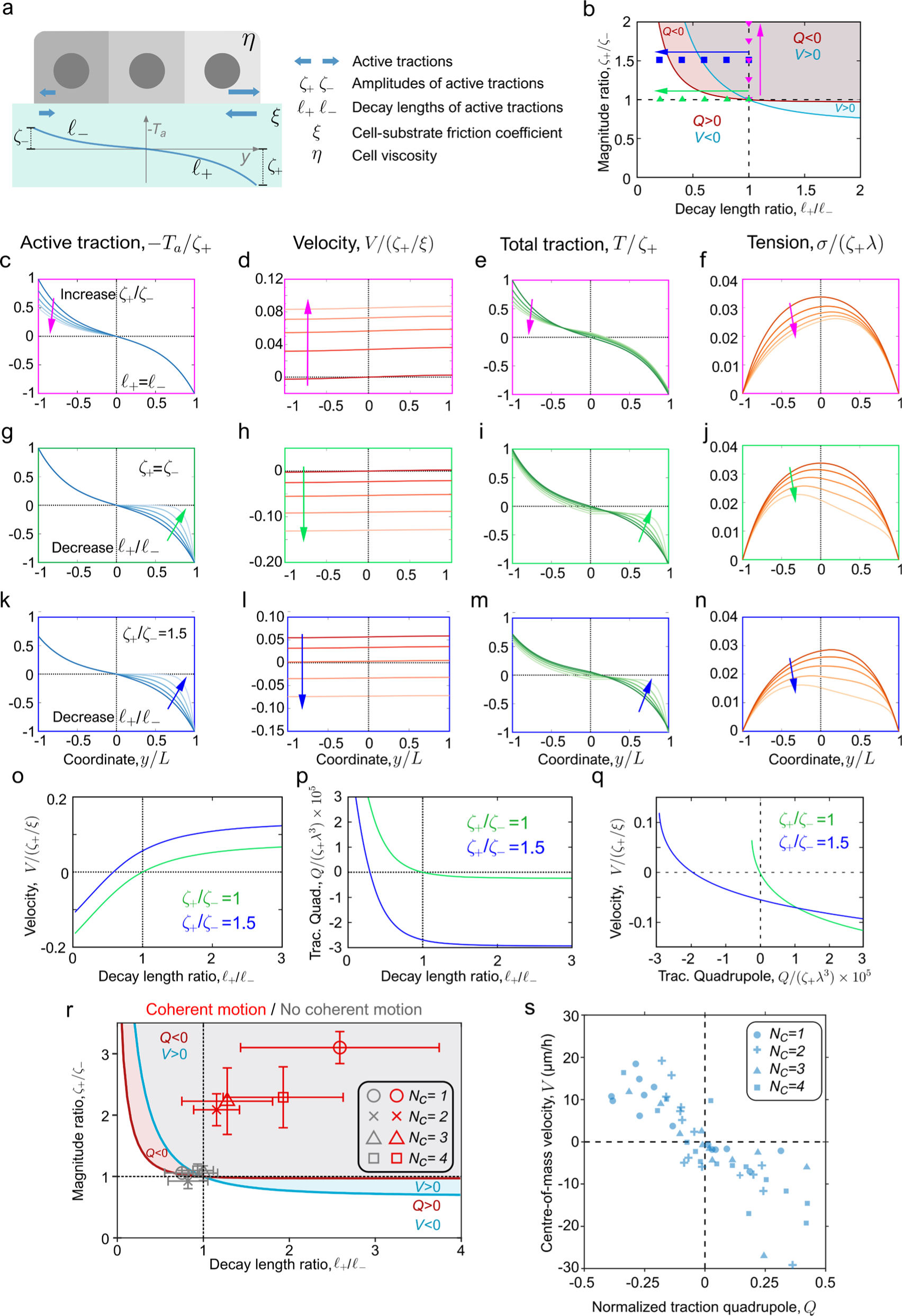
An active polar fluid model explains how traction asymmetries drive migration. **(a)** Scheme of a cell train (grey) that exerts an asymmetric active-traction profile (dark blue curve) on the substrate (light blue). **(b)** Diagram showing the signs of the centre-of-mass velocity *V* and the traction quadrupole *Q* as a function of asymmetries in traction magnitude and decay length. The colour-coded points and arrows correspond to the parameter values used for the profiles in panels c-n. All panels are plotted for *λ*_−_ = 0.4 *L* and *λ* = 10 *L*, with *L* the train length. We chose *λ* ≫ *L* to ensure tension transmission across the entire cell train^52,71^. **(c-n)** Illustrative results of the model in different regimes of asymmetries of the active tractions. Columns correspond to profiles of different quantities. Rows with pink, green, and blue frames correspond to the paths indicated by the corresponding arrows in panel b. **(c-f)** Decreasing active traction magnitude at the left edge (c) results in increased cell velocity (d), as well as in asymmetric total tractions (e) and tension concentrated to the right (f). **(g-j)** Decreasing the decay length of active tractions at the right edge (g) results in negative velocity (h), as well as asymmetric tractions (i) and tension concentrated towards the left (j). **(k-n)** Asymmetry in both the magnitude and the decay length of active tractions. As active tractions become more localized at the right edge (k), velocity decreases in magnitude and eventually changes sign (l). The total tractions (m) and tension (n) profiles also shift from right- to left-concentrated, corresponding to a change of sign of the traction quadrupole *Q*. **(o, p)** Centre-of-mass velocity (o) and traction quadrupole (p) as a function of the decay-length asymmetry of active tractions. Colours correspond to the parameter paths indicated by arrows in panel b. **(q)** Velocity as a function of the quadrupole obtained by varying the decay length ratio. The velocity and the quadrupole change sign simultaneously only when the active-traction magnitude is symmetric (green). **(r)** Diagram as in panel b showing the values of active traction magnitude and decay length ratios obtained from the fits to the experimental profiles in Fig. 3d. Coherently moving cell trains (red points) fall in the region with positive velocity and negative quadrupole (*V* > 0, *Q* < 0), whereas non-coherently moving cell trains (grey points) have symmetric traction profiles that fall close to the origin at (1,1). **(s)** Model simulations recapitulating Fig. 3f. Points are calculated from 60 sets of parameter values ζ_+_, ζ_−_, *l*_+_, *l*_−_, *ξ* drawn from the distributions obtained from fits of the model to all individual cell trains (Methods).

First, we calculated the centre-of-mass velocity 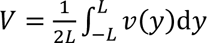. Averaging Eq. (2) over *y* and imposing stress-free boundary conditions [σ(−*L*) = σ(*L*) = 0], we obtained

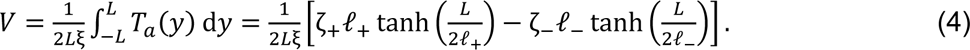

Eq. (4) expresses how motion arises from the balance between the frictional and active contributions to traction. Eq. (4) also shows that the integral of the active traction is the driving force of the motion of cell clusters. This result provides an explicit force-velocity relation for a cell cluster, linking its motion to the asymmetries in the underlying active traction field.

Then, solving Eq. (2) with stress-free boundary conditions as well as velocity and stress continuity at *y* = 0, we obtained the velocity profile *v*(*y*) and used it to calculate the total-traction quadrupole, 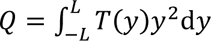, in terms of ζ_+_, ζ_−_, *l*_+_, *l*_−_ and the screening length λ = √η/ξ (Supplementary Note Sections A and B). Fig. 5b summarizes in a 2D diagram how the sign of the velocity and the quadrupole change with the asymmetry ratios ζ_+_/ζ_−_ and *l*_+_/*l*_−_. Below, we illustrate these results by following the paths indicated by the pink, green, and blue arrows. The profiles of all fields at the points along these paths are shown in Fig. 5c-n, and the variation of the velocity and quadrupole along the green and blue paths are plotted in Fig. 5o-q.

First, keeping the decay lengths equal, *l*_+_ = *l*_−_, we varied the ratio between the magnitude of active traction on each edge of the train, ζ_+_/ζ_−_ (pink arrow in Fig. 5b). In the symmetric case ζ_+_/ζ_−_ = 1, there is no possibility for net motion, as the train protrudes with equal force in both directions. To impose an asymmetry, we increase ζ_+_/ζ_−_, making the active traction relatively stronger on the right (Fig. 5c). This imbalance results in cellular motion to the right (positive velocity, Fig. 5d). The profiles of the total traction have an asymmetry analogous to that of the active traction (Fig. 5e), which gives negative quadrupoles *Q* < 0 (Fig. 5b, pink). Accordingly, the tension is concentrated towards the direction of motion (Fig. 5f) and the centre-of-mass velocity is positive and increases with the asymmetry (averages of profiles in Fig. 5d).

Next, we varied the ratio between the decay length of active traction on each side of the train, *l*_+_/*l*_−_, while keeping the magnitudes equal (green arrow in Fig. 5b). Starting again from the symmetric case, we now decreased *l*_+_/*l*_−_, which makes the active traction more localized on the right edge (Fig. 5g). Now the total traction and tension profiles shift towards the left, which gives a positive quadrupole *Q* > 0 and drives leftwards migration, *V* < 0 (Fig. 5h-j and Fig. 5b, green).

In the two cases presented above, *V* has the opposite sign of *Q* and its magnitude increases with that of *Q* (green lines in Fig. 5o-q), consistent with our experimental results (Fig. 3e,f). Our theory also predicts additional scenarios in which the two sources of spatial asymmetries (magnitude vs localization) in active traction have competing effects on cell motion (Supplementary Note Section C). In these cases, there are parameter regions in which *V* and *Q* have the same sign (blue path in Fig. 5b, illustrated in Fig. 5k-n), even if the magnitude of *V* always increases with that of *Q* (blue curves in Fig. 5o-q).

Our model so far assumed that the friction coefficient was uniform. To assess the sensitivity of our theoretical results on this assumption, we generalized the model to account for a non-uniform friction coefficient with the same profile as the active traction (Supplementary Note Section D), which could result from a non-uniform distribution of focal adhesions^54^. The velocity, tension, and traction profiles are only slightly modified with respect to the case with uniform friction coefficient (Extended Data Fig. 5), showing that our conclusions also hold in the presence of a non-uniform friction coefficient.

### Fitting the theory to the experiments

Having analysed the different scenarios that the theory predicts, we fitted the model to the experimental averaged traction profiles in Fig. 3d, and we extracted the values of the active traction parameters ζ_+_, ζ_−_, *l*_+_, *l*_−_ and the friction term *ξV* (Extended Data Fig. 6). This analysis allowed us to position the cell trains of different lengths in the model parameter space (Fig. 5r). As expected, cell trains with no coherent motion fall at the origin of the diagram (1,1), reflecting the symmetry in traction profiles. By contrast, cell trains displaying coherent motion fall in the top right-hand quadrant (*Q*<0 and *V*>0), indicating an asymmetry in both the traction amplitude and the decay length.

Finally, we used our model to recapitulate the experimental relationship between traction quadrupole and velocity (Fig. 3f). We fitted the traction profiles of all individual cell trains and obtained distributions of the parameter values ζ_+_, ζ_−_, *l*_+_, *l*_−_, *ξ*. From these distributions we generated a set of 60 simulated cell trains. For each of these simulated trains, we calculated the traction quadrupole *Q* and used Eq. (4) to compute *V*. We then plotted *V* against *Q* in Fig. 5s and obtained a correlation analogous to the experimental relationship between ⟨*V*⟩_*t*_ and 〈*Q*〉_*t*_ in Fig. 3f.

Overall, our theory provides a force-velocity relation for collective cell migration (Eq. (4)) and captures the link between motion and the strength of the mechanical asymmetries in cell clusters quantified by the traction quadrupole (Fig. 5q). Beyond capturing our experimental measurements (Figs. 3f and 5r, s), our theory reveals that the velocity-quadrupole relation depends on the interplay between asymmetries in the magnitude and localization of cellular forces, providing quantitative predictions for future experiments.

## Discussion

In this work we studied the features that allow cell groups to undergo coherent migration. By using optogenetics to generate leader cells “on demand” we tested the hypothesis that a highly protrusive cell at the edge of a group suffices to guide its global motion. We ruled out this hypothesis: while inducing protrusive activity of one cell is sufficient to direct its own migration and that of one follower, it is insufficient to direct the migration of larger groups. Through direct measurement of cellular forces and theoretical modelling, we showed that collective cell migration takes place when there is a global asymmetry in the tension profile within the cluster. Thus, a leader can only lead if followers provide such an opportunity by contributing to establish an asymmetric mechanical pattern.

The relation between cellular forces and velocities is one of the most fundamental and unresolved problems in cell migration^46,55–59^. Because inertia and viscous drag against the surrounding fluid are negligible, the sum of tractions exerted by cells on the substrate adds up to zero at all times and is therefore not indicative of neither the magnitude nor the direction of cell velocity. Previous work at the single-cell level established that asymmetries in the spatial distribution of tractions correlate with the direction in which a cell moves^43,45^. However, a general quantitative relationship between force and velocity was lacking both at the single and collective cell levels^45,59,43,60–64^. Here we showed that cluster velocity increases with stress asymmetry within the cell cluster. To understand the origin of this relationship, we modelled the cluster as a 1D active polar fluid that can generate asymmetric active tractions, either because they display higher magnitude at the leading edge or because they decay faster away from it. This model fits the total traction and tension distributions measured in our experiments, and it reproduces the relationship between tension asymmetry and cell velocity.

Given that an autonomous leader appears to be unable to drag large cohorts of cells, our study raises the question of how leaders and followers communicate, if at all, to organize in the specific asymmetric patterns that enable collective cell migration. Candidate mechanisms include patterns in cell differentiation, paracrine signalling, or intercellular communication across cell junctions^65–70^. In contexts in which collective cell migration is not properly controlled, such as cancer invasion, mechanical asymmetries could be established stochastically, as observed in some of our experiments in which a few groups spontaneously developed a mechanical asymmetry and performed directed migration even in the absence of photoactivation (Fig. 3). By providing the mechanical rules for effective leadership in cell collectives, our study sets an experimental and conceptual basis to test these communication mechanisms systematically.

## Supporting information

Supplementary Figures and Theory note

## Methods

### Cloning

The TIAM–CRY2-mCherry plasmid was constructed as detailed previously for lentiviral vectors ^37^. The CIBN-GFP-CAAX plasmid was a gift from Chandra Tucker (Denver, Colorado, United States)^40^.

### Cell culture

MDCK strain II cells were cultured in minimum essential medium with Earle’s Salts and l-glutamine (Gibco) supplemented with 10% v/v foetal bovine serum (FBS; Gibco), 100 U ml penicillin and 100 μg/ml streptomycin. Cells were maintained at 37 °C in a humidified atmosphere with 5% CO2. Opto-MDCK-Rac1 fluorescent stable cell lines were obtained by lentiviral transduction of CIBN-GFP-CAAX and TIAM-CRY2-mCherry and two rounds of flow cytometry-based sorting.

### Polyacrylamide gels

We prepared polyacrylamide gels with a stiffness of 18kPa according to a previously established protocol and functionalized them with Sulpho-SANPAH (Thermo Fisher Scientific)^72^. For the gels we prepared a solution of 0.16% bis-acrylamide and 7.5% acrylamide, 0.01% v/v 200-nm-diameter dark-red fluorescence carboxylate-modified beads (Fluospheres, Thermo Fisher Scientific), 0.5% v/v ammonium persulfate (Sigma Aldrich) and 0.05% tetramethylethylenediamine (Sigma Aldrich), in PBS. We placed a 22ul drop of unpolymerized gel on a glass-bottom MatTek 35 mm dish and immediately covered it with an 18mm circular coverslip. The gels were then allowed to polymerize at room temperature for 1 hour and then covered with PBS before removing the circular coverslip. Functionalization of the gel surface was achieved by incubation with a solution of 2 mg/ml Sulpho-SANPAH under ultraviolet light for 7 minutes (wavelength of 365 nm at a distance of 5 cm). Then, two washes of PBS were performed for 2.5 minutes under mild agitation to remove excess Sulpho-SANPAH. The gels were then immediately used for microcontact printing of fibronectin lines.

### Microcontact Printing

Stamps for microcontact printing of 20 μm lines were fabricated from SU8-50 masters that had been raised using conventional photolithography. For the 50 μm lines, the masters were produced by polymerizing a thin layer (20 μm) of photopolymerizing resin (NOA61, Norland) using a UV photopatterning device (PRIMO, Alvéole) coupled to an inverted microscope (Ti Eclipse, Nikon). In both cases the masters contained tens of identical parallel line patterns ∼10 mm x 5 mm. Within each pattern the lines were spaced 80 µm from each other. Uncured Polydimethylsiloxane (PDMS, Sylgard, Dow Corning) was poured on the masters and cured overnight at 65 °C. Solid PDMS stamps were then cut-out and peeled off from the master and their patterned surfaces were treated with a 15 s discharge from a handheld corona surface treater (APS-CD-20AC, Aurora Pro Scientific). Immediately following this, they were covered with a 100 µl drop of a solution of 20 µg/ml fibronectin (fibronectin from human plasma, Sigma Aldrich) and 15 µg/ml fibrinogen-Alexa488 conjugate (F13191, Thermo Fisher) and incubated at room temperature for 1 h. After this, excess incubating solution was removed, and stamps were dried with a nitrogen gun. A polyacrylamide gel was dried thoroughly with a nitrogen gun and the stamp was laid on top of it, with the patterned face in contact with the gel surface. Gel and stamp were left in contact for 1 h after which 1 ml of PBS was added to the MatTek dish. After 1 h the stamp was lifted, and the patterned gels were passivated by incubating overnight at 4 °C with a solution of 0.1 mg/ml PLL-g-PEG in PBS. Finally, the passivating solution was removed, and the gels were covered with a 300 µl drop of PBS and were immediately used for cell-seeding.

### Cell seeding

Opto-MDCK-Rac1 cells were detached from their culture flask using trypsin and resuspended in culture medium. A 300ul drop containing 2×10^4^ cells was placed on a micropatterned gel that had been sterilized under UV light in a cell-culture hood for 15 min. Cells were allowed to adhere for 1 h before unattached cells were removed by a gentle wash with warm cell culture media. The samples were then left to incubate for 12h in 2ml cell media containing thymidine 2 mM (T9250243 1G, Sigma) at 37 °C in a humidified atmosphere with 5% CO_2_.

### Photoactivation experiments and fluorescence imaging

Experiments were carried out on a Zeiss LSM880 confocal microscope running the software Zeiss ZEN2.3 SP1 FP3 (black, version 14.0.24.201), and using a Plan Apochromat 20X 0.8 NA objective. Regions of the sample containing cell groups (trains or clusters) of interest were identified through the eyepieces using white light illumination with a long pass red filter (cut off at 630 nm). Only cell groups that were isolated from other cells on the same line were used for the experiments. The sample was rotated in the plane of the microscope stage so that the lines of the micropatterns appeared vertical in images. Integrity of the micropatterned lines in each region was verified by acquiring a single image with a low-intensity 488nm laser scan, visualizing the signal of the Fibrinogen-Alexa488 conjugate present in the protein-coating. Regions with discontinuous or broken patterns were not used for experiments. After this step, 45 minutes were let pass to allow for the unbinding of CRY2/CIBN and for any induced activation of Rac1 to return to basal levels^39^. Following this, the imaging was started. Three channels were acquired: 561 nm to excite mCherry, 633 nm to excite the fluorescent microspheres and 633 nm transmitted light to obtain a bright-field image. Scanning was performed with a pixel-size of 0.17 µm and a pixel dwell time of 0.35 µs. Up to four fields of view were acquired, with imaging occurring every 3 minutes in a multiposition timelapse. At each position the fluorescence autofocus algorithm of ZEN was run using the microsphere fluorescence as reference. Initially, a baseline phase of 2 h with no photoactivation was acquired. Following this, a subgroup of cell groups was selected to be photoactivated and a rectangular illumination region was drawn on the free edge of one of their edges using the ROI tool of ZEN. Imaging was then resumed as before, and during every subsequent imaging acquisition the photoactivation regions were scanned with the 488 nm laser and the same pixel dwell time as before. The imaging and photoactivation continued for 4 to 5 h with a frequency of 3 minutes. Every 5 image acquisitions (i.e., every 15 min) the positions of the photoactivating regions were manually adjusted according to the movement of the targeted cells. This was done to keep the region of induced lamellipodia at the same relative position within the cell group. This operation required less than 2 minutes. At the end of each experiment, cells were detached from the gel using Versene 1X (Life Technologies) and a reference image of the fluorescent beads was acquired for TFM calculations^42^. Cell groups that merged or touched other cells, and cell groups containing cells that divided during the experiment were not considered for analysis.

### Image analysis

For all timepoints, mCherry fluorescence images were semi-automatically segmented using a custom written MATLAB script and FIJI. A first segmentation was obtained based on the triangle thresholding algorithm, and then any mistakes were manually corrected. The binary masks thus obtained were used to measure cell group dynamics (edge trajectories and centre of mass trajectory). In the photoactivated cases, the images were reoriented (if necessary) so that the photoactivation ROI was at a positive *y* distance from the centre of the cell group. Edge trajectories were smoothed by adjacent averaging with a span of 5 points. Lamellipodium growth was calculated by first dividing the train segmentation in two by bisecting its major axis. The average area (*A*) of the top half (i.e., the half subject to photoactivation) was calculated in the hour prior to and following photoactivation (*A*^∗^). Lamellipodium growth was defined as the difference between these two areas: Δ*A* = *A*^∗^ − *A*.

### Quantifying directed motion

Motility of an edge was characterized using the set of its instantaneous velocities. For each trajectory a two-sided Wilcoxon signed rank test was applied to assess if the set of instantaneous velocities was significantly different from a set with null median. If it was not, the edge was considered to be not significantly motile in any direction. In the case of non-photoactivated trains this approach was applied to trajectories lasting for the whole duration of the experiment, while in the other cases it was applied separately to the trajectories before and during photoactivation, but in order to rule out any transient effect caused by photoactivation we left out the first hour after its application. For a cell train, directed and antidirected motion were defined as the cases when the aforementioned analysis yielded that the median velocities of both edges were significantly non-zero and of the same sign.

### Traction force microscopy and traction force data analysis

All traction computations and the following analyses of traction forces were carried out with custom-written MATLAB scripts. Fourier transform traction microscopy was used to measure traction forces^29,44,73^. The displacement fields of the fluorescence microspheres were obtained using a home-made particle imaging velocimetry algorithm (PIV) using square interrogation windows of side 40 pixels with an overlap of 0.8. The segmented binary masks of the cell trains obtained from the mCherry fluorescence were used to segment the tractions for each train at each time point. The axial profiles of the tractions were calculated by averaging the y-component of the tractions, *T*_*y*_, across the width of the segmented cell trains, at every time point, yielding 〈*T*_*y*_〉_*x*_. The axial lengths of these profiles were normalized to unit length for each time point and the tractions were averaged together in groups according to the train’s behaviour (directed/antidirected motion, or not). In the photoactivated cases, only the timepoints starting 1 hour after photoactivation were considered. The same averaging procedure was applied to obtain axial tension profiles from 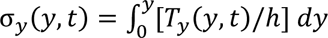, where ℎ is the height of the cell and is approximated to be 10 µm. The normalized second moment of the traction field was calculated as *Q*(*t*) = (∫ 〈*T*_*y*_〉_*x*_*y*^2^dy)/ (∫|〈*T*_*y*_〉_*x*_|*y*^2^ δψ), where *y* is the spatial coordinate relative to the centre of mass, and was then averaged over time, adapting the durations to the photoactivated and non-photoactivated cases as explained above.

### Calculation of cell-cell tension

Calculations were performed under the assumption that the cell train behaves as a 1D material. In that case the Cauchy stress tensor is reduced to a scalar, *σ*, which we refer to it as tension. The equation of mechanical equilibrium in 1D reads: 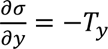, hence internal tension can be obtained by direct integration of the traction field^29,74,75^.

### Kymographs

Kymographs of tractions were obtained by averaging across the *x* axis the 2D traction maps of *T*_*y*_(*y*, *t*), at each individual timepoint. The values of 〈*T*_*y*_⟩_*x*_ were plotted in a colour-coded figure in the order and spacing given by the experiment’s acquisition times.

### Immunostainings

Opto-MDCK-Rac1 cells were photoactivated for 15 min as explained above and immediately fixed with paraformaldehyde 4% during 15 min at room temperature (RT). Then, cells were washed three times with PBS. The immunostainings were performed at RT using tris-buffered saline (TBS) containing 1.6% (v/v) fish gelatine (G7765, Merck) as the basal buffer. First, permeabilization was carried out by treating the samples with 0.1% Triton X-100 (T8787, Sigma-Aldrich) during 45 min. After that, samples were incubated for 90 min with the primary antibody (rabbit anti-phospho-paxillin; 69363s, Cell Signaling) diluted 1:100. After three 3 min washes, samples were incubated for 90 min with the secondary antibody (Alexa Fluor-647 goat anti-rabbit; A-21245, ThermoFisher) diluted 1:200. Finally, they were washed four times for 3 minutes with PBS and mounted in Mowiol reagent (81381, Merck). The image acquisition was done on a Zeiss LSM880 confocal microscope running the software Zeiss ZEN2.3 SP1 FP3 (black, version 14.0.24.201), and using a Plan Apochromat 20X 0.8 NA objective. The photoactivated cells were identified for imaging by using the microscope software to return to the previously stored positions and confirming visually that the same field-of-view had been reached.

### Fits of the theoretical model to the experimental results

Eq. 1 was fitted to the experimental data displayed in the top row of Fig. 3d. *T*_*a*_ is given by Eq. 2 in terms of *l*_+_, *l*_−_, ζ_+_, ζ_−_ and *L*. As the velocity *v* varies weakly in space, it was replaced in Eq. 1 with the average velocity *V* for simplicity. Thus, the fits yield values for the parameters ζ_+_, ζ_−_, *l*_+_, *l*_−_, *ξV*. The fit was performed using the MATLAB function *fitnlm*.

### Simulating the relationship between Q and V

To generate the simulated plot of *Q* against *V* (Fig. 4s) we fitted the model equations to experimental traction profiles 〈*T*_y_⟩_*x*_ as described above, but for each coherently moving cell train. This yielded a set of fit parameters (ζ_+_, ζ_−_, *l*_+_, *l*_−_, *ξV*) for each of the 60 moving trains. We used each train’s center of mass velocity *V* to obtain values of *ξ* and calculated its average (*ξ*⟩ for each value of *N*_*c*_. We did not consider the fits that had diverging values of *l*_+_ and *λ*_−_, leaving us with 43 out of 60 fits. We pooled these into empirical distributions of ζ_+_, ζ_−_, *l*_+_, *l*_−_, one for each value of train length. We then characterized the mean and standard deviation of each of these distributions. To generate the plot, we then randomly extracted 15 sets of parameters ζ_+_, ζ_−_, *l*_+_, *l*_−_ from Gaussian distributions with the same mean and standard deviation as the empirical distributions. In this way we obtained 60 sets of input parameters (using (*ξ*⟩ for the corresponding *N*_*c*_) whose distribution closely resemble the empirical values. For each of these 60 model configurations, we have calculated the net active traction force according to Eq. 3 and the quadrupole of the total traction force as described above, resulting in the scatter plot in Fig. 4s.

### Statistical tests, box plots and Sankey diagrams

All statistical significance analysis was performed using a two-sided Wilcoxon rank sum test, as implemented by the MATLAB functions ranksum and signrank. Box plots in all figures show the median value, and the 25^th^ and 75^th^ percentiles of the data. The whiskers have a set length of 1.5 times the interquartile range (difference between 75^th^ and 25^th^ percentile). Sankey diagrams were generated using the custom MATLAB function *Sankey flow chart*^76^.

## Acknowledgments

We thank all the members of our groups for their discussions and support. We thank Susana Usieto and Mònica Purciolas for technical assistance. We also thank Chandra Tucker and Simon De Beco for sharing plasmids used in this work. We thank Pascal Silberzan for sharing experimental data with us. Finally, we thank Ion Andreu, Marija Matejčić and Amy Beedle for discussions and feedback on the manuscript. This paper was funded by the Generalitat de Catalunya (AGAUR SGR-2017-01602 to X.T., the CERCA Programme, and “ICREA Academia” awards to P.R-C.); the Spanish Ministry for Science and Innovation MICCINN/FEDER (PID2021-128635NB-I00, MCIN/AEI/ 10.13039/501100011033 and “ERDF-EU A way of making Europe” to X.T.,, PID2019-110298GB-I00 to P.R-C.); European Research Council (101097753 to P.R-C. and Adv-883739 to X.T.); Fundació la Marató de TV3 (project 201903-30-31-32 to X.T.); European Commission (H2020-FETPROACT-01-2016-731957 to P.R-C. and X.T.); the European Union’s Horizon 2020 research and innovation programme (under the Marie Skłodowska Curie Grant agreement ID: 796883 to L.R.); La Caixa Foundation (LCF/PR/HR20/52400004 to P.R-C. and X.T.); IBEC is recipient of a Severo Ochoa Award of Excellence from the MINECO.

## Author contributions

L.R., L.V. and X.T. conceived the project. L.V. designed and performed preliminary experiments. L.R. and J.F.A. designed and performed experiments. L.R. and S.G. analysed data. J.F.A. and P.R-C. contributed technical expertise, materials and discussion. R.A. developed the model. L.R., R.A. and X.T. wrote the manuscript. All authors revised the completed manuscript.

## Competing interests

The authors declare no competing financial interests.

## Code availability

Analysis procedures and code implementing the model are available from the corresponding authors on reasonable request.

## Data availability

The data that support the findings of this study are available from the corresponding authors on reasonable request. Extended Data is available for this paper. Correspondence and requests for materials should be addressed to L.R., R.A. or X.T.

## Notes

### Competing Interest Statement

The authors have declared no competing interest.

## References

1. Schaller, V., Weber, C., Semmrich, C., Frey, E. & Bausch, A. R. Polar patterns of driven filaments. Nature 467, 73–77 (2010).

2. Boudet, J. F. et al. From collections of independent, mindless robots to flexible, mobile, and directional superstructures. Sci. Robot. 6, eabd0272 (2021).

3. Silverberg, J. L., Bierbaum, M., Sethna, J. P. & Cohen, I. Collective Motion of Humans in Mosh and Circle Pits at Heavy Metal Concerts. Phys. Rev. Lett. 110, 228701 (2013).

4. Suzuki, R., Weber, C. A., Frey, E. & Bausch, A. R. Polar pattern formation in driven filament systems requires non-binary particle collisions. Nat. Phys. 11, 839–843 (2015).

5. Ben-Jacob, E., Cohen, I. & Levine, H. Cooperative self-organization of microorganisms. Adv. Phys. 49, 395–554 (2000).

6. Gómez-Nava, L., Bon, R. & Peruani, F. Intermittent collective motion in sheep results from alternating the role of leader and follower. Nat. Phys. 18, 1494–1501 (2022).

7. Yllanes, D., Leoni, M. & Marchetti, M. C. How many dissenters does it take to disorder a flock? New J. Phys. 19, 103026 (2017).

8. Pearce, D. J. G. & Giomi, L. Linear response to leadership, effective temperature, and decision making in flocks. Phys. Rev. E 94, 022612 (2016).

9. Pinkoviezky, I., Couzin, I. D. & Gov, N. S. Collective conflict resolution in groups on the move. Phys. Rev. E 97, 032304 (2018).

10. Couzin, I. D., Krause, J., Franks, N. R. & Levin, S. A. Effective leadership and decision-making in animal groups on the move. Nature 433, 513–516 (2005).

11. Nagy, M., Ákos, Z., Biro, D. & Vicsek, T. Hierarchical group dynamics in pigeon flocks. Nature 464, 890–893 (2010).

12. Omelchenko, T., Vasiliev, J. M., Gelfand, I. M., Feder, H. H. & Bonder, E. M. Rho-dependent formation of epithelial “leader” cells during wound healing. Proc. Natl. Acad. Sci. 100, 10788–10793 (2003).

13. Poujade, M. et al. Collective migration of an epithelial monolayer in response to a model wound. Proc. Natl. Acad. Sci. 104, 15988–15993 (2007).

14. Khalil, A. A. & Friedl, P. Determinants of leader cells in collective cell migration. Integr. Biol. 2, 568 (2010).

15. Mayor, R. & Etienne-Manneville, S. The front and rear of collective cell migration. Nat. Rev. Mol. Cell Biol. 17, 97–109 (2016).

16. Theveneau, E. & Linker, C. Leaders in collective migration: are front cells really endowed with a particular set of skills? F1000Research 6, 1899 (2017).

17. Yang, Y. & Levine, H. Leader-cell-driven epithelial sheet fingering. Phys. Biol. 17, 046003 (2020).

18. Pinheiro, D., Kardos, R., Hannezo, É. & Heisenberg, C.-P. Morphogen gradient orchestrates pattern-preserving tissue morphogenesis via motility-driven unjamming. Nat. Phys. 18, 1482–1493 (2022).

19. Camley, B. A. & Rappel, W.-J. Physical models of collective cell motility: from cell to tissue. J. Phys. Appl. Phys. 50, 113002 (2017).

20. Martinson, W. D. et al. Dynamic fibronectin assembly and remodeling by leader neural crest cells prevents jamming in collective cell migration. eLife 12, e83792 (2023).

21. Kozyrska, K. et al. p53 directs leader cell behavior, migration, and clearance during epithelial repair. Science 375, eabl8876 (2022).

22. Yamaguchi, N., Mizutani, T., Kawabata, K. & Haga, H. Leader cells regulate collective cell migration via Rac activation in the downstream signaling of integrin β1 and PI3K. Sci. Rep. 5, 7656 (2015).

23. Reffay, M. et al. Interplay of RhoA and mechanical forces in collective cell migration driven by leader cells. Nat. Cell Biol. 16, 217–223 (2014).

24. Hino, N. et al. A feedback loop between lamellipodial extension and HGF-ERK signaling specifies leader cells during collective cell migration. Dev. Cell 57, 2290–2304.e7 (2022).

25. Vishwakarma, M. et al. Mechanical interactions among followers determine the emergence of leaders in migrating epithelial cell collectives. Nat. Commun. 9, 3469 (2018).

26. Law, R. A. et al. Cytokinesis machinery promotes cell dissociation from collectively migrating strands in confinement. Sci. Adv. 9, eabq6480 (2023).

27. Cai, D. et al. Mechanical Feedback through E-Cadherin Promotes Direction Sensing during Collective Cell Migration. Cell 157, 1146–1159 (2014).

28. Vishwakarma, M., Spatz, J. P. & Das, T. Mechanobiology of leader–follower dynamics in epithelial cell migration. Curr. Opin. Cell Biol. 66, 97–103 (2020).

29. Trepat, X. et al. Physical forces during collective cell migration. Nat. Phys. 5, 426–430 (2009).

30. Caussinus, E., Colombelli, J. & Affolter, M. Tip-Cell Migration Controls Stalk-Cell Intercalation during Drosophila Tracheal Tube Elongation. Curr. Biol. 18, 1727–1734 (2008).

31. Arima, S. et al. Angiogenic morphogenesis driven by dynamic and heterogeneous collective endothelial cell movement. Development 138, 4763–4776 (2011).

32. Wang, X., He, L., Wu, Y. I., Hahn, K. M. & Montell, D. J. Light-mediated activation reveals a key role for Rac in collective guidance of cell movement in vivo. Nat. Cell Biol. 12, 591–597 (2010).

33. Labernadie, A. et al. A mechanically active heterotypic E-cadherin/N-cadherin adhesion enables fibroblasts to drive cancer cell invasion. Nat. Cell Biol. 19, 224–237 (2017).

34. Cheung, K. J., Gabrielson, E., Werb, Z. & Ewald, A. J. Collective Invasion in Breast Cancer Requires a Conserved Basal Epithelial Program. Cell 155, 1639–1651 (2013).

35. Vilchez Mercedes, S. A., et al. Decoding leader cells in collective cancer invasion. Nat. Rev. Cancer 21, 592–604 (2021).

36. Machacek, M. et al. Coordination of Rho GTPase activities during cell protrusion. Nature 461, 99–103 (2009).

37. de Beco, S. et al. Optogenetic dissection of Rac1 and Cdc42 gradient shaping. Nat. Commun. 9, 4816 (2018).

38. Drozdowski, O. M., Ziebert, F. & Schwarz, U. S. Optogenetic control of migration of contractile cells predicted by an active gel model. Commun. Phys. 6, 1–12 (2023).

39. Valon, L. et al. Predictive Spatiotemporal Manipulation of Signaling Perturbations Using Optogenetics. Biophys. J. 109, 1785–1797 (2015).

40. Kennedy, M. J. et al. Rapid blue-light–mediated induction of protein interactions in living cells. Nat. Methods 7, 973–975 (2010).

41. Valon, L., Marín-Llauradó, A., Wyatt, T., Charras, G. & Trepat, X. Optogenetic control of cellular forces and mechanotransduction. Nat. Commun. 8, 14396 (2017).

42. Roca-Cusachs, P., Conte, V. & Trepat, X. Quantifying forces in cell biology. Nat. Cell Biol. 19, 742–751 (2017).

43. Hennig, K. et al. Stick-slip dynamics of cell adhesion triggers spontaneous symmetry breaking and directional migration of mesenchymal cells on one-dimensional lines. Sci. Adv. 6, eaau5670 (2020).

44. Butler, J. P., Tolić-Nørrelykke, I. M., Fabry, B. & Fredberg, J. J. Traction fields, moments, and strain energy that cells exert on their surroundings. Am. J. Physiol.-Cell Physiol. 282, C595–C605 (2002).

45. Tanimoto, H. & Sano, M. A Simple Force-Motion Relation for Migrating Cells Revealed by Multipole Analysis of Traction Stress. Biophys. J. 106, 16–25 (2014).

46. Delanoë-Ayari, H., Rieu, J. P. & Sano, M. 4D Traction Force Microscopy Reveals Asymmetric Cortical Forces in Migrating Dictyostelium Cells. Phys. Rev. Lett. 105, 248103 (2010).

47. Costa, G. et al. Asymmetric division coordinates collective cell migration in angiogenesis. Nat. Cell Biol. 18, 1292–1301 (2016).

48. Hayashi, S. & Dong, B. Shape and geometry control of the Drosophila tracheal tubule. Dev. Growth Differ. 59, 4–11 (2017).

49. Weigelin, B., Bakker, G.-J. & Friedl, P. Intravital third harmonic generation microscopy of collective melanoma cell invasion. IntraVital 1, 32–43 (2012).

50. Alert, R. & Trepat, X. Physical Models of Collective Cell Migration. Annu. Rev. Condens. Matter Phys. 11, 77–101 (2020).

51. Pérez-González, C. et al. Active wetting of epithelial tissues. Nat. Phys. 15, 79–88 (2019).

52. Alert, R., Blanch-Mercader, C. & Casademunt, J. Active Fingering Instability in Tissue Spreading. Phys. Rev. Lett. 122, 088104 (2019).

53. Blanch-Mercader, C. et al. Effective viscosity and dynamics of spreading epithelia: a solvable model. Soft Matter 13, 1235–1243 (2017).

54. Delanoë-Ayari, H., Bouchonville, N., Courçon, M. & Nicolas, A. Linear Correlation between Active and Resistive Stresses Provides Information on Force Generation and Stress Transmission in Adherent Cells. Phys. Rev. Lett. 129, 098101 (2022).

55. Brückner, D. B. et al. Stochastic nonlinear dynamics of confined cell migration in two-state systems. Nat. Phys. 15, 595–601 (2019).

56. Chan, C. E. & Odde, D. J. Traction Dynamics of Filopodia on Compliant Substrates. Science 322, 1687–1691 (2008).

57. Bangasser, B. L. et al. Shifting the optimal stiffness for cell migration. Nat. Commun. 8, 15313 (2017).

58. Bergert, M. et al. Force transmission during adhesion-independent migration. Nat. Cell Biol. 17, 524–529 (2015).

59. Sakamoto, R., Izri, Z., Shimamoto, Y., Miyazaki, M. & Maeda, Y. T. Geometric trade-off between contractile force and viscous drag determines the actomyosin-based motility of a cell-sized droplet. Proc. Natl. Acad. Sci. 119, e2121147119 (2022).

60. Godeau, A. L. et al. 3D single cell migration driven by temporal correlation between oscillating force dipoles. eLife 11, e71032 (2022).

61. Carlsson, A. E. Mechanisms of cell propulsion by active stresses. New J. Phys. 13, 073009 (2011).

62. Amiri, B., Heyn, J. C. J., Schreiber, C., Rädler, J. O. & Falcke, M. On multistability and constitutive relations of cell motion on Fibronectin lanes. Biophys. J. (2023) doi:10.1016/j.bpj.2023.02.001.

63. Basan, M., Elgeti, J., Hannezo, E., Rappel, W.-J. & Levine, H. Alignment of cellular motility forces with tissue flow as a mechanism for efficient wound healing. Proc. Natl. Acad. Sci. 110, 2452–2459 (2013).

64. Ron, J. E. et al. Polarization and motility of one-dimensional multi-cellular trains. Biophys. J. 122, 4598–4613 (2023).

65. Camley, B. A. Collective gradient sensing and chemotaxis: modeling and recent developments. J. Phys. Condens. Matter 30, 223001 (2018).

66. Ruppel, A. et al. Force propagation between epithelial cells depends on active coupling and mechano-structural polarization. eLife 12, e83588 (2023).

67. George, M., Bullo, F. & Campàs, O. Connecting individual to collective cell migration. Sci. Rep. 7, 9720 (2017).

68. Zimmermann, J., Camley, B. A., Rappel, W.-J. & Levine, H. Contact inhibition of locomotion determines cell–cell and cell–substrate forces in tissues. Proc. Natl. Acad. Sci. 113, 2660–2665 (2016).

69. Boutillon, A. et al. Guidance by followers ensures long-range coordination of cell migration through α-catenin mechanoperception. Dev. Cell 57, 1529–1544.e5 (2022).

70. Campanale, J. P. & Montell, D. J. Who’s really in charge: Diverse follower cell behaviors in collective cell migration. Curr. Opin. Cell Biol. 81, 102160 (2023).

71. Alert, R. & Casademunt, J. Role of Substrate Stiffness in Tissue Spreading: Wetting Transition and Tissue Durotaxis. Langmuir 35, 7571–7577 (2019).

## Methods References

72. Serra-Picamal, X. et al. Mechanical waves during tissue expansion. Nat. Phys. 8, 628–634 (2012).

73. Serra-Picamal, X., Conte, V., Sunyer, R., Muñoz, J. J. & Trepat, X. Mapping forces and kinematics during collective cell migration. in Methods in Cell Biology vol. 125 309–330 (Elsevier, 2015).

74. Tambe, D. T. et al. Collective cell guidance by cooperative intercellular forces. Nat. Mater. 10, 469–475 (2011).

75. Tambe, D. T. et al. Monolayer Stress Microscopy: Limitations, Artifacts, and Accuracy of Recovered Intercellular Stresses. PLOS ONE 8, e55172 (2013).

76. Borau, C. Sankey flow chart. https://www.mathworks.com/matlabcentral/fileexchange/101516-sankey-flow-chart (2022)

