## Supplementary Figures and Theory note for "Optogenetic generation of leader cells reveals a force-velocity relation for collective cell migration"

**Supplementary Movie 1 –** Effect of photoactivation on a cell train with $N_c = 1$. Composite of brightfield and CIBN-GFP-CAAX fluorescence excited at 488 nm. Scale bar 20 μm.

**Supplementary Movie 2 -** Effect of photoactivation on a cell train with $N_c = 2$. Composite of brightfield and CIBN-GFP-CAAX fluorescence excited at 488 nm. Scale bar 20 μm.

**Supplementary Movie 3 -** Effect of photoactivation on a cell train with $N_c = 3$. Composite of brightfield and CIBN-GFP-CAAX fluorescence excited at 488 nm. Scale bar 20 μm.

**Supplementary Movie 4 -** Effect of photoactivation on a cell train with $N_c = 4$. Composite of brightfield and CIBN-GFP-CAAX fluorescence excited at 488 nm. Scale bar 20 μm.

**Supplementary Movie 5 -** Effect of photoactivation on a cell island. Composite of brightfield and CIBN-GFP-CAAX fluorescence excited at 488 nm. Scale bar 20 μm.

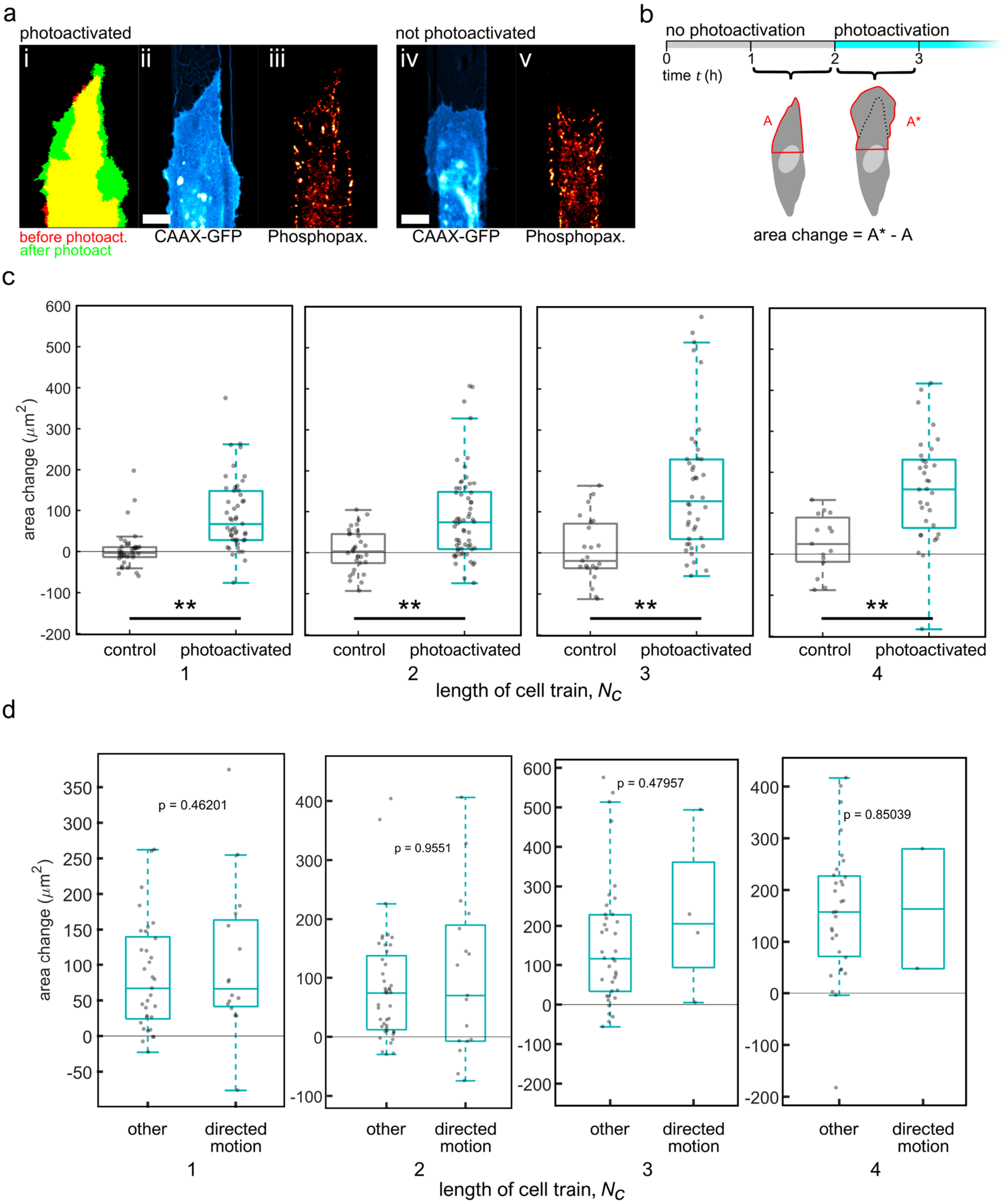


**Extended Data Figure 1. Photoactivation induces lamellipodium growth. (a)** (i) Segmentation of a lamellipodium before and after photoactivation, (ii) membrane fluorescence of lamellipodium after photoactivation, (iii) focal adhesions after photoactivation, (iv) a naturally occurring lamellipodium, (v) focal adhesions in the naturally occurring lamellipodium (scale bar 10 µm). **(b)** Scheme of how lamellipodium area is calculated. The area of the top half of the cell is averaged during 1 h prior to (A) and following (A*) photoactivation (these same time intervals are considered also for control cells). The change of lamellipodium size is calculated as the difference between A* and A. **(c)** Lamellipodium area for trains of different lengths, comparing control cases and photoactivated trains. Photoactivation induces lamellipodium growth in nearly all cases. Statistical significance quantified by a two-sided Wilcoxon rank sum test, ** indicates $p<0.01$. **(d)** Lamellipodium growth of photoactivated trains undergoing directed motion compared with other trains. Lamellipodium growth is not significantly different between these two subpopulations. Statistical significance quantified by a two-sided Wilcoxon rank sum test.

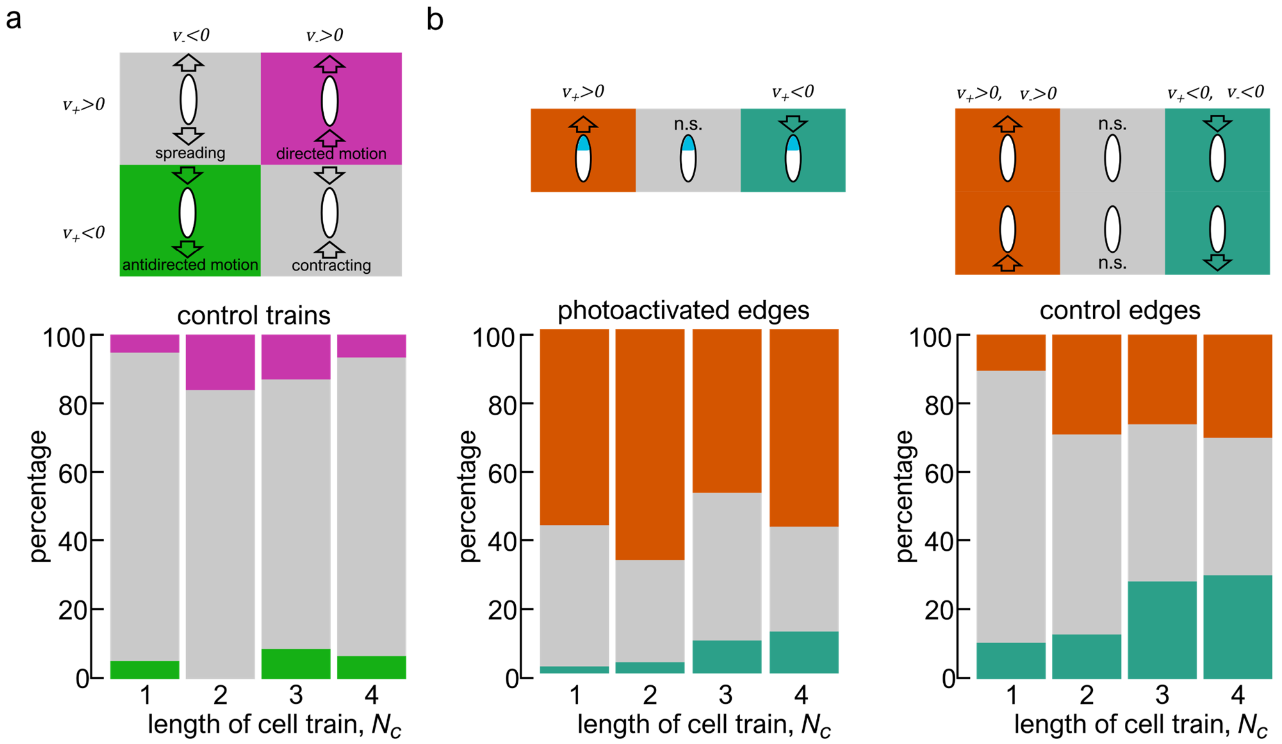


**Extended Data Figure 2. Cell train and edge motion. (a)** Directed migration in control trains. In absence of photoactivation there is an equal probability for upwards (directed) or downwards (antidirected) migration. **(b)** Effect of photoactivation on edge motion, compared with control cases. The bars show the percentage of photoactivated edges that have significant velocities in the direction of photoactivation (magenta), in the opposing direction (green), or that have non-significant velocities (grey). For all values of $N_c$ more than 50% of the cell trains have edge velocities biased in the direction of the induced lamellipodium. In the control case, both directions are equally probable. For increasing values of $N_c$ the photoactivated edges total sample sizes are n=49, n=61, n=42, n=33, respectively, and for the control edges total sample sizes are n=38, n=31, n=23, n=15.

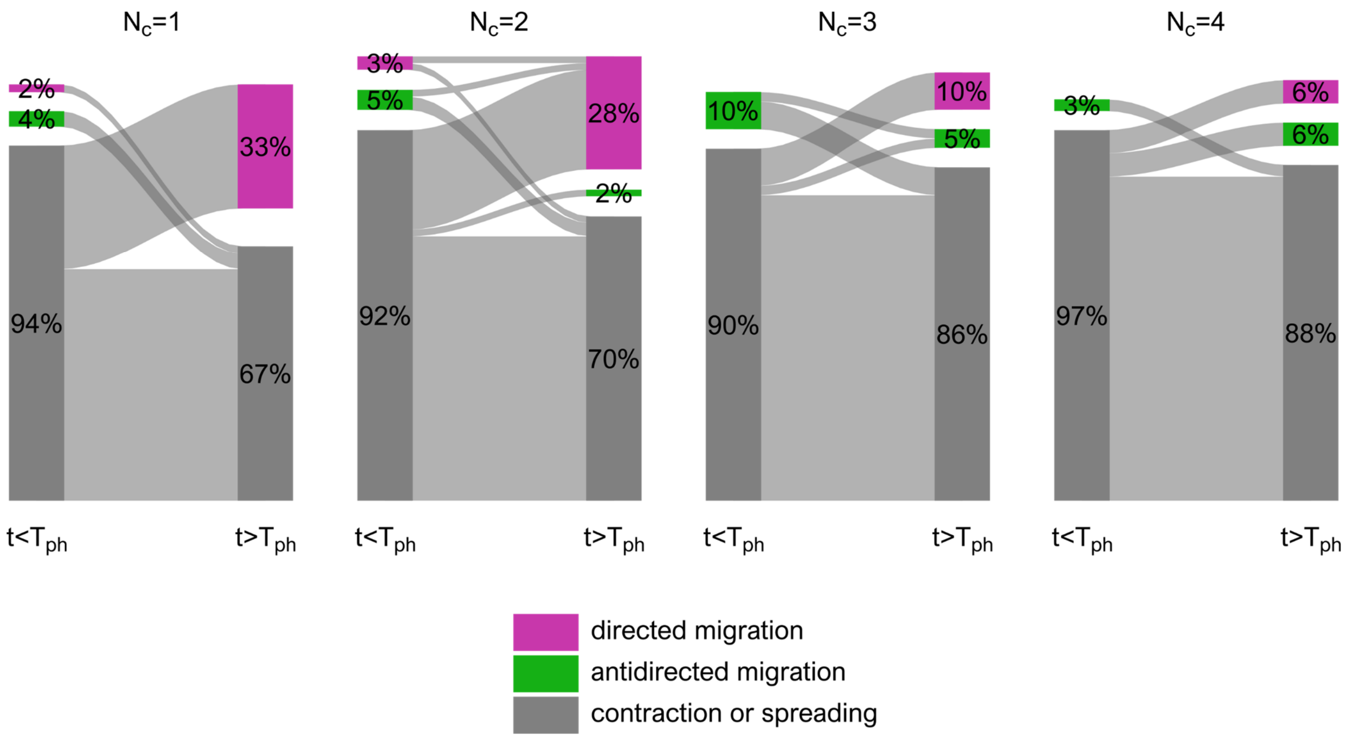


**Extended Data Figure 3. Photoactivated cell trains that undergo directed migration are not undergoing directed migration prior to photoactivation.** Motion of cell trains of different lengths before and during photoactivation. For each value of $N_c$ the column on the left shows the type of migration of the trains that will be photoactivated, while the column on the right shows the type of migration of the same trains during photoactivation (same data as Fig. 2e). The stripes connecting the columns show how cell trains changed their migration type due to photoactivation. $T_{ph}$ stands for the time at which photoactivation starts and $t$ is time.

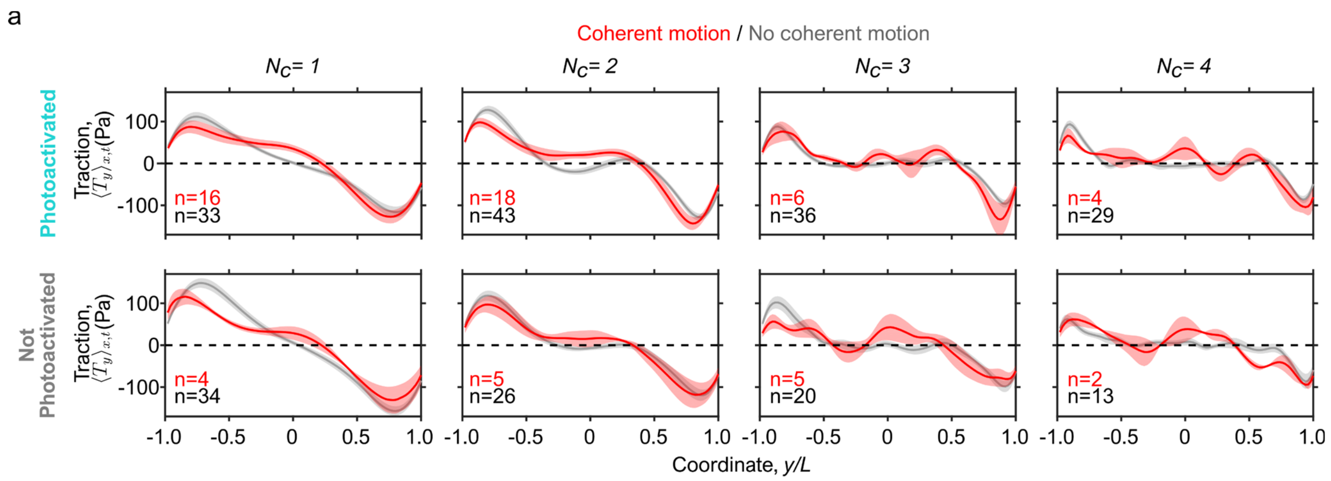


**Extended Data Figure 4. Effect of photoactivation on traction forces. (a)** Average profiles of the longitudinal component of the traction for cell trains undergoing coherent motion (red) and for other trains (grey). The top row shows only photoactivated trains, the bottom row non-photoactivated trains. Shaded regions along the curves show the standard error of the mean. Averages are over different cell trains; number of cell trains $n$ is indicated on the plot.

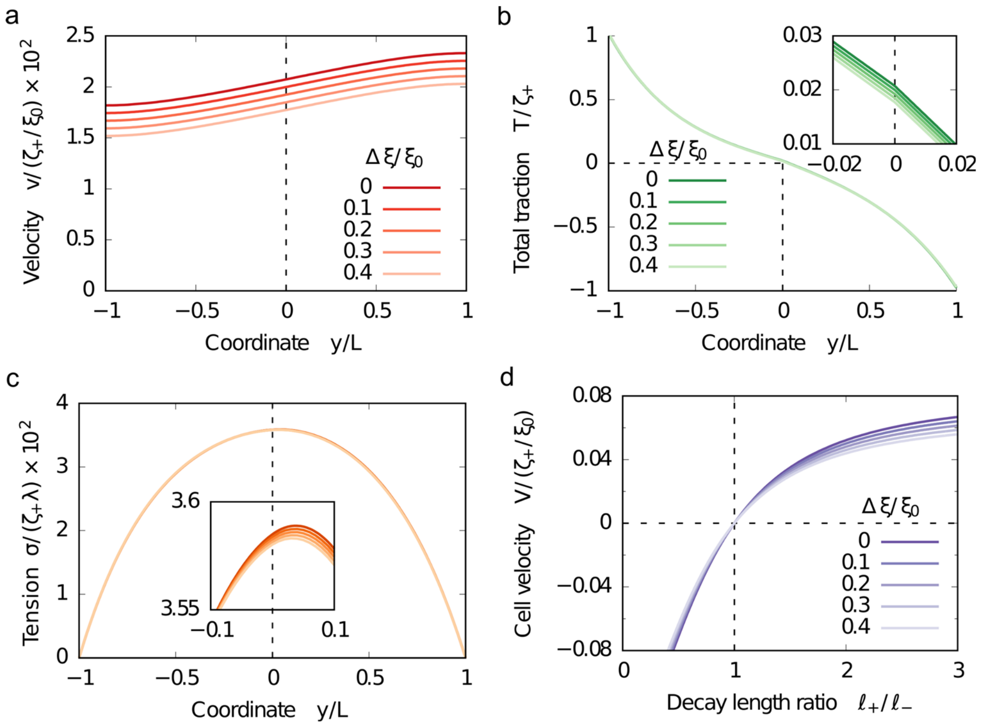


**Extended Data Figure 5. Role of non-uniform friction on train motion. (a-c)** Predicted profiles of velocity (a), total traction (b), and tension (c) for different values of the relative strength of friction non-uniformity, $\Delta\xi/\xi_0$, varied from 0% to 40%. Insets help to visualize the small effect of friction non-uniformity. The theory with non-uniform friction is explained in Section D of the Supplementary Note. **(d)** Centre-of-mass velocity as a function of the decay-length asymmetry of the active tractions, as in Fig. 4o, shown for different values of the relative friction non-uniformity. The effects of friction non-uniformity are small in all cases. In all panels, we chose $\zeta_+ = \zeta_-$, $\ell_- = 0.4\,L$, and $\lambda = 10\,L$. In panels a-c, we chose $\ell_+/\ell_- = 1.25$.

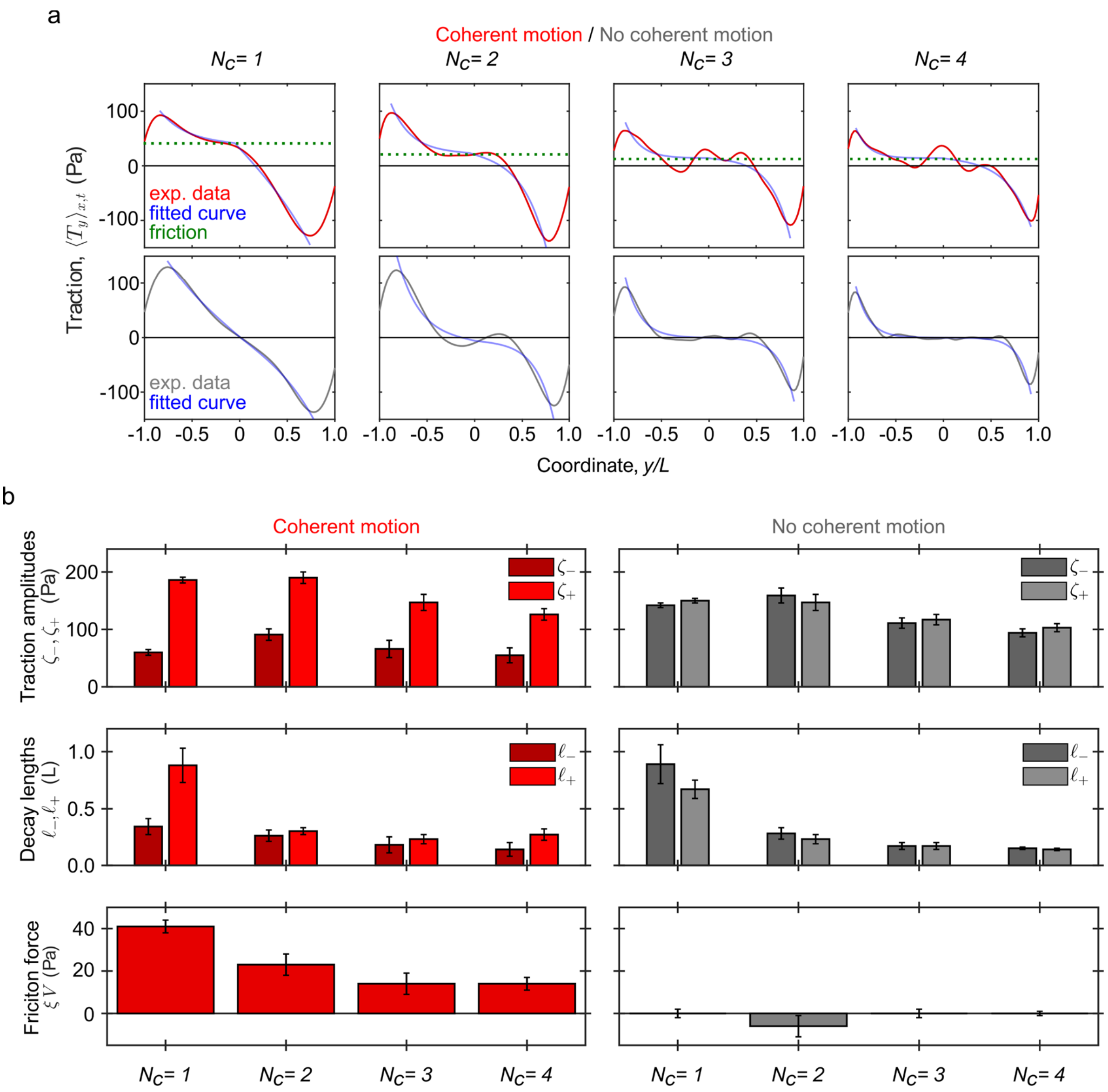


**Extended Data Figure 6. Fits of the predicted traction profiles to the experimental data. (a)** Model (blue curves) and experimental (red and grey curves) average longitudinal tractions for cell trains undergoing coherent motion (top row, red curves) and for other trains (bottom row, grey curves). In the top row, dashed green lines show the level of friction force, as also reported in the bottom left plot of panel (b). **(b)** Parameter values of the model (Eq. (1)) obtained from the fits.

**Supplementary Table 1**

Values of the statistical significance in the main figures and supplementary figures.

Fig. 2c, p-values

| $N_c$ | photoactivated | control | comparison |
|---|---|---|---|
| 1 | 0.00007 | 0.16170 | 0.00005 |
| 2 | 0.00000 | 0.10380 | 0.00490 |
| 3 | 0.06890 | 0.31550 | 0.03070 |
| 4 | 0.24910 | 0.76150 | 0.37360 |

Fig. 3e, p-values

| $N_c$ | |
|---|---|
| 1 | 0.010 |
| 2 | 0.003 |
| 3 | 0.025 |
| 4 | 0.022 |

Extended Data Fig 1c, p-values

| $N_c$ | |
|---|---|
| 1 | 1.0E-08 |
| 2 | 7.4E-05 |
| 3 | 2.0E-06 |
| 4 | 2.2E-04 |

### Supplementary Note for
### "Optogenetic generation of leader cells reveals a force-velocity relation for collective cell migration"


Leone Rossetti,[1,*] Steffen Grosser,[1] Juan Francisco Abenza,[1,2] Léo Valon,[3]
Pere Roca-Cusachs,[1,4] Ricard Alert,[5,6,7,†] and Xavier Trepat[1,2,4,8,‡]

[1]*Institute for Bioengineering of Catalonia (IBEC), The Barcelona Institute for Science and Technology (BIST), 08028 Barcelona, Spain*
[2]*Centro de Investigación Biomédica en Red en Bioingeniería, Biomateriales y Nanomedicina (CIBER-BBN), 08028 Barcelona, Spain*
[3]*Department of Developmental and Stem Cell Biology, Institut Pasteur, CNRS UMR 3738, 75015 Paris, France*
[4]*Facultat de Medicina, Universitat de Barcelona, 08036 Barcelona, Spain*
[5]*Max Planck Institute for the Physics of Complex Systems, 01187 Dresden, Germany*
[6]*Center for Systems Biology Dresden, 01307 Dresden, Germany*
[7]*Cluster of Excellence Physics of Life, TU Dresden, 01062 Dresden, Germany*
[8]*Institució Catalana de Recerca i Estudis Avançats (ICREA), Barcelona, Spain*


(Dated: January 17, 2024)


In this Supplementary Note, we provide details of our theory. In Section A, we detail the calculation of the velocity, tension, and traction profiles shown in Fig. 5c-n. The result for the center-of-mass velocity is given directly in Eq. 4 of the Main Text. In Section B, we calculate the three lowest multipole moments of the traction field: the monopole, dipole, and quadrupole. In Section C, we discuss the relation between these traction multipoles and motion. Our discussion in the Main Text (Fig. 5b,o-r) focuses on the quadrupole, which quantifies the asymmetry of the traction field. Finally, in Section D, we generalize the model to include an inhomogeneous friction coefficient.

#### A. Velocity, tension, and traction profiles

To predict the profiles of velocity, tension, and traction, we solve Eq. 2 of the Main Text:

$$v(y) = \begin{cases} V_1^+ \sinh(y/\lambda) + V_2^+ \cosh(y/\lambda) + \frac{\zeta_+/\xi}{1-(\lambda/\ell_+)^2} \frac{\sinh(y/\ell_+)}{\sinh(L/\ell_+)}; & y \geq 0 \\ V_1^- \sinh(y/\lambda) + V_2^- \cosh(y/\lambda) + \frac{\zeta_-/\xi}{1-(\lambda/\ell_-)^2} \frac{\sinh(y/\ell_-)}{\sinh(L/\ell_-)}; & y < 0, \end{cases} \tag{S1}$$

where $\lambda \equiv \sqrt{\eta/\xi}$ is the screening length, which characterizes the extent to which substrate friction screens the transmission of tension across the cell train. To determine the integration constants $V_1^+$, $V_2^+$, $V_1^-$, and $V_2^-$, we impose stress-free boundary conditions on both cell edges, $y = \pm L$, as well as continuity of both tension and velocity at the central point $y = 0$ that defines the switch in the piecewise active traction profile (Eq. 3 and Fig. 5a). Altogether, we have

$$\sigma_+(L) = 0, \qquad \sigma_-(-L) = 0, \qquad \sigma_+(0) = \sigma_-(0), \qquad v_+(0) = v_-(0), \tag{S2}$$

where the $\pm$ subscripts indicate the solutions for $y \geq 0$ and $y < 0$ sides of the train, respectively. From these conditions, we obtain

$$V_1^+ = \frac{1}{2\xi\cosh(L/\lambda)}\left[-\frac{\zeta_+\lambda/\ell_+}{1-\lambda^2/\ell_+^2}\frac{\cosh(L/\lambda)+\cosh(L/\ell_+)}{\sinh(L/\ell_+)} + \frac{\zeta_-\lambda/\ell_-}{1-\lambda^2/\ell_-^2}\frac{\cosh(L/\lambda)-\cosh(L/\ell_-)}{\sinh(L/\ell_-)}\right], \tag{S3a}$$

$$V_1^- = \frac{1}{2\xi\cosh(L/\lambda)}\left[\frac{\zeta_+\lambda/\ell_+}{1-\lambda^2/\ell_+^2}\frac{\cosh(L/\lambda)-\cosh(L/\ell_+)}{\sinh(L/\ell_+)} - \frac{\zeta_-\lambda/\ell_-}{1-\lambda^2/\ell_-^2}\frac{\cosh(L/\lambda)+\cosh(L/\ell_-)}{\sinh(L/\ell_-)}\right], \tag{S3b}$$

$$V_2^+ = V_2^- = \frac{1}{2\xi\sinh(L/\lambda)}\left[\frac{\zeta_+\lambda/\ell_+}{1-\lambda^2/\ell_+^2}\frac{\cosh(L/\lambda)-\cosh(L/\ell_+)}{\sinh(L/\ell_+)} - \frac{\zeta_-\lambda/\ell_-}{1-\lambda^2/\ell_-^2}\frac{\cosh(L/\lambda)-\cosh(L/\ell_-)}{\sinh(L/\ell_-)}\right]. \tag{S3c}$$

These solutions give the velocity profiles, and they also determine the tension and total traction profiles as $\sigma(y) = \eta v'(y)$ and $T(y) = \xi v(y) - T_{\text{a}}(y)$, respectively. Figure 5 shows these profiles for different combinations of active traction asymmetries.


*
†
‡

##### B. Traction multipole moments

Here, we compute the multipole moments of the traction field. First, the monopole moment is the net, integrated traction force, and it must vanish because cell motion occurs in almost inertia-free conditions:

$$M \equiv \int_{-L}^{L} T(y)\,\mathrm{d}y = 0. \tag{S4}$$

This condition is identical to the tension continuity condition. This equivalence can be seen by integrating the force balance equation, $\partial_y \sigma = T$, and using the stress-free boundary conditions in Eq. (S2):

$$M = \int_{-L}^{L} T(y)\,\mathrm{d}y = \int_{-L}^{L} \partial_y \sigma(y)\,\mathrm{d}y = \sigma_+(L) - \sigma_+(0) + \sigma_-(0) - \sigma_-(-L) = 0. \tag{S5}$$

Using the total traction profiles $T(y)$ obtained as explained in Section A, we obtain the dipole and quadrupole moments as

$$D \equiv \int_{-L}^{L} T(y)\,y\,\mathrm{d}y = \frac{\zeta_+\lambda^2}{1-\lambda^2/\ell_+^2}\left[\frac{\lambda}{\ell_+}\frac{\tanh(L/\lambda)}{\tanh(L/\ell_+)} - 1\right] + \frac{\zeta_-\lambda^2}{1-\lambda^2/\ell_-^2}\left[\frac{\lambda}{\ell_-}\frac{\tanh(L/\lambda)}{\tanh(L/\ell_-)} - 1\right], \tag{S6}$$

$$\begin{aligned} Q \equiv& \int_{-L}^{L} T(y)\,y^2\,\mathrm{d}y \\ =& \frac{2\zeta_+\lambda^2\ell_+}{1-\lambda^2/\ell_+^2}\left\{\left[1-\frac{\lambda^2}{\ell_+^2}\right]\tanh\left(\frac{L}{2\ell_+}\right) + \frac{L\lambda}{\ell_+^2}\left[\coth(L/\lambda)\coth(L/\ell_+) - \operatorname{csch}(L/\lambda)\operatorname{csch}(L/\ell_+)\right] - \frac{L}{\ell_+}\right\} \\ &- \frac{2\zeta_-\lambda^2\ell_-}{1-\lambda^2/\ell_-^2}\left\{\left[1-\frac{\lambda^2}{\ell_-^2}\right]\tanh\left(\frac{L}{2\ell_-}\right) + \frac{L\lambda}{\ell_-^2}\left[\coth(L/\lambda)\coth(L/\ell_-) - \operatorname{csch}(L/\lambda)\operatorname{csch}(L/\ell_-)\right] - \frac{L}{\ell_-}\right\}. \end{aligned} \tag{S7}$$

The sign of the dipole moment $D$ indicates whether tractions point inward or outward on average. As cells exert contractile traction patterns on the substrate, we imposed active tractions $-T_\mathrm{a}$ that point inward, toward the interior of the cell train (Eq. 3 and Fig. 5a). Accordingly, the dipole moment is negative ($D < 0$).

Respectively, the sign of the quadrupole moment $Q$ indicates fore-aft asymmetries of the traction field, placing a stronger weight on larger distances from the center, equivalently to a moment of inertia. If tractions are shifted towards $y > 0$, the quadrupole is negative ($Q < 0$); if tractions are shifted towards $y < 0$, the quadrupole is positive ($Q > 0$). In the Main Text and below, we discuss how the different asymmetries of the traction field either cooperate or compete to set the traction quadrupole, and how the quadrupole is related to the velocity of a cell cluster.

##### C. Relation between traction multipoles and motion

How are the traction multipole moments related to motion? Because inertia is negligible at the cellular and multicellular scales, the net force is zero. Cells perform force-free motion. Therefore, the traction monopole is always zero.

The dipole is not directly related to the center-of-mass velocity $V$ of a cell cluster. For example, symmetric traction profiles have non-zero dipole moment but do not drive cell motion, giving a vanishing velocity $V = 0$. Asymmetric traction profiles also have a non-zero dipole moment, but they drive motion. Put in a different way, two cell clusters can have the same dipole moment but one can be static and the other one moving.

Here, through both theory and experiments, we find that the quadrupole is more directly related to cell velocity. For completely symmetric traction profiles, both quadrupole and velocity vanish. In general, asymmetries in the active traction profile give both a non-zero quadrupole and velocity. Both theoretically and experimentally, we find that stronger asymmetries, quantified by larger quadrupole magnitudes, correspond to higher velocities (Fig. 3f and Fig. 5q,s). Our work therefore establishes a direct quantitative relation between motion and traction asymmetries in collective cell migration. Thus, our results explain and generalize previous observations of correlations between the traction multipole and symmetry breaking in single-cell motion[1–3].

Beyond capturing our experimental findings, our theory also predicts additional scenarios that unveil the complexity of the velocity-quadrupole relation. Strikingly, we find that the quadrupole does not necessarily vanish when the cell velocity does (blue curve in Fig. 5q). In other words, there are specific asymmetric traction profiles for which the cell velocity vanishes. Therefore, observing a static cell cluster does not imply that its traction quadrupole is zero. Conversely, measuring a non-zero quadrupole does not imply that the cell or cluster must move.

In general terms, our theoretical results establish a non-trivial relation between cell motion and the traction quadrupole (Fig. 5b,q). This relation is non-trivial because both quantities depend on the interplay between two sources of asymmetry: the magnitude $\zeta$ and the decay length $\ell$ of cellular active tractions (Eq. 3). As a result, the relation between velocity $V$ and traction quadrupole $Q$ is not as simple as could be expected: $V$ and $Q$ are not forced to have opposite signs (Fig. 5b,o-q).

In addition to depending on the active traction parameters, the traction quadrupole picks a dependence on the cell or cluster viscosity $\eta$ via $\lambda = \sqrt{\eta/\xi}$ (Eq. (S7)). This dependence comes from the contribution of cell-substrate friction $\xi v$ to the traction tension, $T = \xi v - T_{\text{a}}$. By contrast, the center-of-mass velocity is independent of the viscosity $\eta$ (Eq. 4). Therefore, changing the viscosity affects the traction quadrupole but leaves the net velocity unaffected, which further emphasizes that cell or cluster motion does not have a simple one-to-one relation with the traction quadrupole.

In our model, the situations in which the velocity-quadrupole relation is not straightforward can be understood as follows. Cell motion is driven by the integral of the active traction profile (Eq. 4). Asymmetries in traction magnitude and decay length can either add up or cancel each other out. For example, a stronger but more localized traction on the right can be compensated or even overpowered by a weaker but more spread traction on the left, giving zero or even leftward motion. In contrast, as it is analogous to a moment of inertia, the traction quadrupole weighs tractions applied further from the center of mass much more than tractions applied closer to the center of mass. Therefore, tractions more localized at the cell edge can easily dominate the sign of the quadrupole, even if the active traction integral, and hence cell motion, is dominated by more spread tractions on the other side of the cell or cluster.

To illustrate this competition, we fixed the asymmetry in magnitude, setting $\zeta_+/\zeta_- = 1.5$, and we varied the asymmetry in decay lengths (blue arrow in Fig. 5b). When the decay lengths are equal, $\ell_+/\ell_- = 1$, cells move to the right ($V > 0$; Fig. 5o, blue), driven by the stronger active traction on the right ($Q < 0$; Fig. 5p, blue). However, as the ratio $\ell_+/\ell_-$ decreases, active tractions become more localized on the right (Fig. 5k), and the velocity decreases (Fig. 5l). Eventually, asymmetries in magnitude and localization compensate each other and motion stops (point with $V = 0$ in Fig. 5o, blue). In this case, the active traction, total traction, and tension profiles are still asymmetric (Fig. 5k-n), but the train remains static ($V = 0$). As tractions become even more localized, the direction of motion is reversed ($V < 0$; Fig. 5l and Fig. 5o, blue), but the tension remains biased rightwards ($Q < 0$; Fig. 5n and Fig. 5p, blue). At even higher localization, the total traction and tension profiles shift from being biased rightwards to leftwards (Fig. 5m-n), and the quadrupole finally changes sign ($Q > 0$; Fig. 5p, blue). These results show that cell trains can be static even if the traction profile is asymmetric.

##### D. Non-uniform friction

In our model so far, for simplicity, we assumed a uniform cell-substrate friction coefficient $\xi$. Cell-substrate friction is due in part to non-specific interactions, not mediated by focal adhesions. We expect this contribution to be rather uniform throughout the cell-substrate interface, and hence our choice of a uniform friction coefficient. However, cell-substrate friction can also have a contribution from specific interactions through focal adhesions. Focal adhesions typically accumulate towards the edges of cell trains and layers (see, for example, Fig. 2k in Ref.[4]). This non-uniform distribution of focal adhesions contributes to active tractions being stronger at the edges, and it can similarly make friction stronger at the edges. Here, we incorporate this effect in our theory by taking a space-dependent friction coefficient $\xi(y)$. Treating this effect perturbatively, we show that, for our choice of parameter values, the velocity, tension, and traction profiles are only slightly modified with respect to the case with uniform friction (Extended Data Fig. 5). Hence, our conclusions hold as well in the presence of non-uniform friction.

To include non-uniform friction, we express the friction coefficient as

$$\xi(y) = \xi_0 + \delta\xi(y), \tag{S8}$$

where $\xi_0$ represents the uniform contribution from non-specific interactions, and $\delta\xi(y)$ represents the non-uniform contribution from specific focal adhesions. Since focal adhesions transmit both active and passive forces across the cell-substrate interface, we take the same spatial dependence for $\delta\xi(y)$ as for the active tractions $T_{\text{a}}(y)$ (Eq. 3 in the Main Text). Thus,

$$\delta\xi(y) = \begin{cases} \Delta\xi_+ \frac{\sinh(y/\ell_+)}{\sinh(L/\ell_+)}; & y \geq 0 \\ \\ -\Delta\xi_- \frac{\sinh(y/\ell_-)}{\sinh(L/\ell_-)}; & y < 0. \end{cases} \tag{S9}$$

Whereas active tractions point in opposite directions at the two edges of a cell train, the friction coefficient increases towards both edges; hence the minus sign in front of the positive coefficient $\Delta\xi_-$.

With non-uniform friction, the force balance (Eq. 2 in the Main Text) becomes

$$\eta v''(y) = \xi(y)v(y) - T_{\text{a}}(y). \tag{S10}$$

To solve it analytically, we treat the effects of friction non-uniformity perturbatively. Thus, we call $v_0(y)$ the solution to the uniform-friction case (Eq. (S1)), and we expand the velocity field as $v(y) = v_0(y) + \delta v(y)$, where $\delta v(y)$ is the perturbation due to friction non-uniformity. Introducing it into Eq. (S10) and expanding to linear order in the perturbations, we obtain

$$\eta\delta v''(y) = \xi_0\delta v(y) + \delta\xi(y)v_0(y). \tag{S11}$$

The full solution is of the form

$$\delta v_\pm(y) = U_1^\pm \sinh(y/\lambda) + U_2^\pm \cosh(y/\lambda) + A_\pm \sinh(y/\ell_\pm)\sinh(y/\lambda) + B_\pm \sinh(y/\ell_\pm)\cosh(y/\lambda) \\ + C_\pm \sinh^2(y/\ell_\pm) + D_\pm \cosh^2(y/\ell_\pm) + E_\pm \cosh(y/\ell_\pm)\sinh(y/\lambda) + F_\pm \cosh(x/\ell_\pm)\cosh(y/\lambda), \quad \text{(S12)}$$

where the subscripts $\pm$ indicate the solutions for the $y \geq 0$ and $y < 0$ sides of the train, respectively. Here, the constants $A$ to $F$ are obtained from the particular solution of the inhomogeneous term:

$$A_\pm = \pm\frac{\Delta\xi_\pm}{\xi_0}\frac{1}{(\lambda/\ell_\pm)^2 - 4}\frac{V_1^\pm}{\sinh(L/\ell_\pm)}, \quad \text{(S13a)}$$

$$B_\pm = \pm\frac{\Delta\xi_\pm}{\xi_0}\frac{1}{(\lambda/\ell_\pm)^2 - 4}\frac{V_2^\pm}{\sinh(L/\ell_\pm)}, \quad \text{(S13b)}$$

$$C_\pm = \mp\frac{\Delta\xi_\pm}{\xi_0}\frac{\zeta_\pm/\xi_0}{1-(\lambda/\ell_\pm)^2}\frac{1-2(\lambda/\ell_\pm)^2}{1-4(\lambda/\ell_\pm)^2}\frac{1}{\sinh^2(L/\ell_\pm)}, \quad \text{(S13c)}$$

$$D_\pm = 2\frac{\lambda^2}{\ell_\pm^2}\frac{C_\pm}{1-2(\lambda/\ell_\pm)^2}, \quad \text{(S13d)}$$

$$E_\pm = -2\frac{\ell_\pm}{\lambda}B_\pm, \quad \text{(S13e)}$$

$$F_\pm = -2\frac{\ell_\pm}{\lambda}A_\pm. \quad \text{(S13f)}$$

Here, $V_1^\pm$ and $V_2^\pm$ are the integration constants of the uniform-friction problem, given in Eq. (S3). Respectively, the integration constants $U_1^\pm$ and $U_2^\pm$ in Eq. (S12) are determined by the boundary conditions in Eq. (S2), which for the perturbations imply

$$\delta\sigma_+(L) = 0, \qquad \delta\sigma_-(-L) = 0, \qquad \delta\sigma_+(0) = \delta\sigma_-(0), \qquad \delta v_+(0) = \delta v_-(0). \quad \text{(S14)}$$

The resulting expressions of $U_1^\pm$ and $U_2^\pm$ are too long to be reproduced here.

Using the solution for the velocity perturbations $\delta v(y)$, we obtain the perturbed profiles of velocity, tension, and total traction. To linear order in perturbations, we have

$$v(y) = v_0(y) + \delta v(y), \quad \text{(S15a)}$$

$$\sigma(y) = \sigma_0(y) + \delta\sigma(y) = \eta v_0'(y) + \eta\delta v'(y), \quad \text{(S15b)}$$

$$T(y) = T_0(y) + \delta T(y) = \xi_0 v_0(y) - T_{\text{a}}(y) + \xi_0\delta v(y) + \delta\xi(y)v_0(y). \quad \text{(S15c)}$$

We show the effect of friction non-uniformity in the case with asymmetric decay lengths, $\ell_+ \neq \ell_-$, but symmetric force strengths: Both active tractions and passive friction have equal magnitudes at both edges, $\zeta_+ = \zeta_-$ and $\Delta\xi_+ = \Delta\xi_- \equiv \Delta\xi$. We then vary the relative strength of friction non-uniformity $\Delta\xi/\xi_0$ from 0 to $40\%$. For our choice of parameter values, motivated by our experimental results, the effect of friction non-uniformity on the velocity, tension, and traction profiles is very small (Extended Data Fig. 5a-c).

Finally, we also study the effect of non-uniform friction on the center-of-mass velocity, defined as $V = \frac{1}{2L}\int_{-L}^{L} v(y)\mathrm{d}y$ as in the Main Text. To this end, we use that $\sigma(y) = \eta v'(y)$ to rewrite the force balance Eq. (S10) as

$$\frac{\sigma'(y)}{\xi(y)} = v(y) - \frac{T_{\text{a}}(y)}{\xi(y)}. \quad \text{(S16)}$$

Integrating over $y$ and using that $\sigma(L) = \sigma(-L) = 0$, we obtain

$$V = \frac{1}{2L}\int_{-L}^{L}\frac{T_{\text{a}}(y)}{\xi(y)}\mathrm{d}y + \frac{1}{2L}\int_{-L}^{L}\frac{\xi'(y)}{\xi^2(y)}\sigma(y)\mathrm{d}y. \quad \text{(S17)}$$

To first order in perturbations, this expression expands into

$$V = V_0 + \delta V = \frac{1}{2L}\int_{-L}^{L}\frac{T_{\text{a}}(y)}{\xi_0}\mathrm{d}y - \frac{1}{2L}\int_{-L}^{L}\frac{T_{\text{a}}(y)}{\xi_0^2}\delta\xi(y)\mathrm{d}y + \frac{1}{2L}\int_{-L}^{L}\frac{\delta\xi'(y)}{\xi_0^2}\eta v_0'(y)\mathrm{d}y. \quad \text{(S18)}$$

This results shows that the perturbation in train velocity $\delta V$, composed of the two last terms, can be computed without the solution for the velocity perturbation profile $\delta v(y)$. The resulting expression is very long and not informative, so we omit it here.

As for the profiles (Extended Data Fig. 5a-c), we vary the relative strength of friction non-uniformity $\Delta\xi/\xi_0$ from $0$ to $40\%$, and we find that its effect on the train velocity is very small (Extended Data Fig. 5d).

Overall, these results show that our conclusions hold as well in the presence of non-uniform friction.


1. Tanimoto, H. & Sano, M. A Simple Force-Motion Relation for Migrating Cells Revealed by Multipole Analysis of Traction Stress. *Biophys. J.* **106**, 16–25 (2014).
2. Hennig, K. *et al.* Stick-slip dynamics of cell adhesion triggers spontaneous symmetry breaking and directional migration of mesenchymal cells on one-dimensional lines. *Sci. Adv.* **6**, eaau5670 (2020).
3. Godeau, A. L. *et al.* 3D single cell migration driven by temporal correlation between oscillating force dipoles. *Elife* **11**, e71032 (2022).
4. Pérez-González, C. *et al.* Active wetting of epithelial tissues. *Nat. Phys.* **15**, 79–88 (2019).
